# Estimates of autozygosity through runs of homozygosity in farmed coho salmon

**DOI:** 10.1101/2020.02.01.930065

**Authors:** Grazyella M. Yoshida, Pablo Cáceres, Rodrigo Marín-Nahuelpi, Ben F. Koop, José M. Yáñez

## Abstract

The characterization of runs of homozygosity (ROH), using high-density single nucleotide polymorphisms (SNPs) allows inferences to be made about the past demographic history of animal populations and the genomic ROH has become a common approach to characterize the inbreeding. We aimed to analyze and characterize ROH patterns and compare different genomic and pedigree-based methods to estimate the inbreeding coefficient in two pure lines (POP A and B) and one recently admixed line (POP C) of coho salmon breeding nuclei, genotyped using a 200K Affymetrix Axiom^®^ myDesign Custom SNP Array. A large number and greater mean length of ROH were found for the two “pure” lines and the recently admixed line (POP C) showed the lowest number and smaller mean length of ROH. The ROH analysis for different length classes suggests that all three coho salmon lines the genome is largely composed of a high number of short segments (<4 Mb), and for POP C no segment >16 Mb was found. A high variable number of ROH, mean length and inbreeding values across chromosomes; positively the consequence of artificial selection. Pedigree-based inbreeding values tended to underestimate genomic-based inbreeding levels, which in turn varied depending on the method used for estimation. The high positive correlations between different genomic-based inbreeding coefficients suggest that they are consistent and may be more accurate than pedigree-based methods, given that they capture information from past and more recent demographic events, even when there are no pedigree records available.

## 1. Introduction

Coho salmon (*Oncorhynchus kisutch*) is one of the six Pacific salmon species which can be found in North America and Asia [1]. In Chile, coho salmon farming began at the end of the 1970s, when about 500,000 eggs were imported from the Kitimat River (British Columbia) and Oregon to start the genetic basis of the Chilean broodstocks [2,3]. The first coho salmon breeding program started in 1992 with rapid growth as the main breeding objective. After four generations of selection for harvest weight, genetic gains of ~10% per generation were reported [2,3].

Genetic improvement programs for aquaculture species have been successfully established for increasing the productivity, for traits like growth and resistance against diseases [4,5]. However, one of the negative consequences of selective breeding is the accumulation of inbreeding, due to the use of related individuals for reproductive purposes [6]. As consequence, a reduction in both the additive genetic variance and diversity is observed, as well as a decrease in the response to selection. Furthermore, inbreeding can result in a phenomenon known as inbreeding depression, defined as a reduction in fitness traits, including growth, survival and reproductive ability, due to the expression of detrimental recessive alleles given the existence of highly homozygous animals in the population [6,7]. Thus, monitoring and managing the inbreeding levels is critical in the operation of genetic improvement programs [8–10].

Inbreeding is traditionally calculated using pedigree records, the estimates might not reflect the true inbreeding level due to 1) the stochastic nature of recombination, 2) the assumption that there are no changes in allele frequencies in time and 3) the persistence of ancestral short segments through time [11]. In addition this approach fails to capture the relatedness among founder animals from the base population [12]. Furthermore, previous studies agreement that errors in pedigrees and incomplete or missing information lead to incorrect or biased inbreeding estimates [13]. The development of genomic technologies, including dense single nucleotide polymorphism (SNP) creates opportunities to estimate inbreeding from genomic-based approaches; for instance, identity by state (IBS) using a genomic relationship matrix [14] or through ROH [15].

ROH are defined as continuous homozygous segments of the individuals’ genome [16], *i.e.*, genomic regions which have identical haplotypes that are identical by state (IBS), which might be a consequence of not random mating or consanguineous mating [17]. Therefore, ROH can be used for quantifying individual autozygosity that occurs when parents have a common ancestor and pass on segments that are identical by decedent (IBD) to the progeny [18]. ROH may provide a more accurate measure of inbreeding levels, compared to using pedigree records [18,19]. Furthermore, the identification and characterization of ROH can provide insights into population history, structure and demographics over time [18,20]. Long ROH segments are indicative of recent IBD, whereas short segments indicate ancient inbreeding, and the sum of all these segments are suggested to be an accurate estimation of the inbreeding level of an individual [21].

Inbreeding studies using genome-wide data were previously reported in humans [16,22,23], cattle [11,19,24–26], swine [27–29], sheep [30], and goats [31]. A recent study reported ROH patterns in rainbow trout populations to show the impact on selection on the genetic diversity in farmed stocks [32]. Studies aimed at ROH pattern characterization and comparisons between coefficients of inbreeding using different approaches are scarce in aquaculture species, due to the necessity of deep and complete pedigree information and dense genomic information. The objectives of this study were: (i) to identify and characterize the ROH patterns in three farmed Chilean coho salmon populations and (ii) to compare estimates of inbreeding coefficients calculated from runs of homozygosity (F_ROH_), genomic relationship matrix (F_GRM_), observed and expected number of homozygous genotypes (F_HOM_), and a pedigree-based relationship matrix (F_PED_).

## 2. Methods

### 2.1 Coho salmon populations and genotypes

Two independent coho salmon populations, managed in two-year reproductive cycles (POP A and POP B) were used in this study and belong to the Pesquera Antares breeding program established in Chile in 1997. Both populations have undergone nine generations of selection for harvest weight, since 1997 and 1998, POPA and B respectively. In addition, POP C is the progeny produced by mating sires from the seventh and dams from eighth generations of POP A and B, respectively. POP C was generated in 2013 to limit inbreeding levels, as suggested by Yáñez et al. [8]. The reproduction system, fish tagging and selection criteria of POP C were described previously [33,34].

Genomic DNA was extracted from fin clips of 88, 45 and 108 animals from POP A, B and C, respectively. The samples were genotyped using a 200K Affymetrix Axiom^®^ myDesign Custom SNP Array developed by the EPIC4 coho salmon genome consortium (http://www.epic-4.org) and built by ThermoFisher Scientific (San Diego, USA). More detail about the array design was previously described by Barria et al. [35]. A genotype quality control was performed in Plink v1.09 [36] using the following parameters to exclude markers: Hardy-Weinberg Equilibrium (HWE) *p-value* < 1e−6, Minor Allele Frequency (MAF) < 0.01 and call rate < 0.90 for genotypes and samples. Furthermore, we retained only the SNP markers that commonly segregated among the three populations.

### 2.2 Principal components and admixture analysis

We used the software Plink v1.09 [36] to evaluate the genetic differentiation among the three coho salmon populations through principal component analysis (PCA). The first two PCAs were plotted using R scripts [37]. The population structure was also examined using a hierarchical Bayesian model implemented in STRUCTURE software v.2.3.4 [38]. We used three replicates of K values ranging from 1 to 12, running of 50,000 iterations and burn-in of 20,000 iterations. To choose the best K value we used the posterior probability values [38].

### 2.3 Runs of homozygosity

Runs of homozygosity analysis was performed separately for all animals in each population using the R package detectRUNS [39]. The following constraints were applied to ROH detected: (i) the minimum number of SNPs included in a ROH was 50, (ii) the minimum length of a ROH was set at 1 Mb, (iii) the maximum distance between adjacent SNPs was 500 Kb, (iv) maximum missing genotypes allowed were 5, (v) density was at least 1 SNP per 50 kb and (vi) sliding windows approach was used to detect ROH for each genotyped animal at each marker position. ROH were classified into five length classes: 1–2, 2–4, 4–8, 8-16 and > 16 Mb, identified as ROH_1–2 Mb_, ROH_2–4 Mb_, ROH_4-8 Mb_, ROH_8–16 Mb_, and ROH_>16 Mb_, respectively.

### 2.4 Inbreeding coefficient

We estimated inbreeding coefficients using three different genomic methods and pedigree relationship matrix (F_PED_). Inbreeding coefficient based on runs of homozygosity (F_ROH_) was estimated for each animal based on all ROHs (ROH_ALL_) and the ROH distribution of five different lengths (ROH_1–2 Mb_, ROH_2–4 Mb_, ROH_4-8 Mb_, ROH_8–16 Mb_, and ROH_>16 Mb_), as follows [40]:

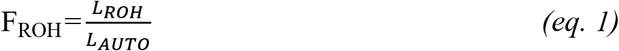

where LROH is the sum of ROH lengths and LAUTO is the total length of genome covered by the genome-wide SNP panel used, assumed to be 1685.79 Mb.

The F_HOM_ was calculated by computing the number of observed and expected homozygous (hom) genotypes for each sample, as follows:

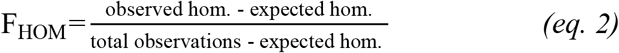

The F_GRM_ was calculated using the genomic relationship matrix (GRM) [14], as follows:

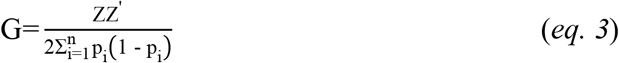

where Z is a genotype matrix that contains the 0 – 2p values for homozygotes, 1 – 2p for heterozygotes, and 2 – 2p for opposite homozygotes, p is the allele frequency of SNP *i*. The diagonal elements of the matrix G represent the relationship of the animal *j* with itself, thus, the genomic inbreeding coefficient is calculated as G_*jj*_ – 1.

Pedigree-based inbreeding coefficients were estimated using the software INBUPGF90 [41]. The pedigree information used was provided by Pesquera Antares breeding program in Chile, for all animals born between 1998 and 2014, 1997 and 2013 and 1998 and 2013 for POP A, B and C respectively.

Pearson correlation between genomic- and pedigree-based inbreeding coefficients was estimated within population using function *cor.test* in R [37].

## 3. Results

### 3.1 Quality control and genomic population structure

The MAF < 0.01 criteria excluded higher numbers of SNPs (~29.9, 19.7 and 18.5 K for POP A, B and C, respectively), and a number of markers between 3.2K to 14.9K were removed to select only common markers segregating across all three populations (Table 1). Thus, out of the initial 135,500 markers, a total of 102,129 markers passed all the QC filtering steps and were shared among the three populations.

**Table 1.** Number of runs of homozygosity (nROH), length (Mb) and standard deviation (SD in Mb) considered all ROH and by classes and for each salmon coho population.

| Class | POP A |  |  | POP B |  |  | POP C |  |  |
| --- | --- | --- | --- | --- | --- | --- | --- | --- | --- |
|  | nROH | Mean length | SD | nROH | Mean length | SD | nROH | Mean length | SD |
| **ROH<sub>ALL</sub>** | 3568 | 5.965 | 7.23 | 1624 | 6.695 | 7.668 | 495 | 3.319 | 3.921 |
| **ROH<sub>1-2 Mb</sub>** | 1165 | 1.284 | 0.389 | 468 | 1.286 | 0.370 | 298 | 1.193 | 0.360 |
| **ROH<sub>2-4 Mb</sub>** | 937 | 2.886 | 0.577 | 400 | 2.831 | 0.530 | 86 | 2.869 | 0.550 |
| **ROH<sub>4-8 Mb</sub>** | 680 | 5.585 | 1.069 | 310 | 5.547 | 1.015 | 74 | 5.337 | 1.170 |
| **ROH<sub>8-16 Mb</sub>** | 463 | 11.308 | 2.323 | 260 | 11.427 | 2.248 | 24 | 11.722 | 1.884 |
| **ROH<sub>>16 Mb</sub>** | 323 | 24.925 | 7.676 | 186 | 23.916 | 8.249 | 13 | 20.443 | 4.056 |

In the PCA analysis, the first two eigenvectors, together, accounted for 29.2% of the total genetic variation and revealed three stratified populations (Figure 1). PCA1 included 22.15% of the total genetic variation and generated the principal clusters to differentiate the three coho salmon populations, whereas PCA2 explained the variation present within each population.

**Figure 1.**
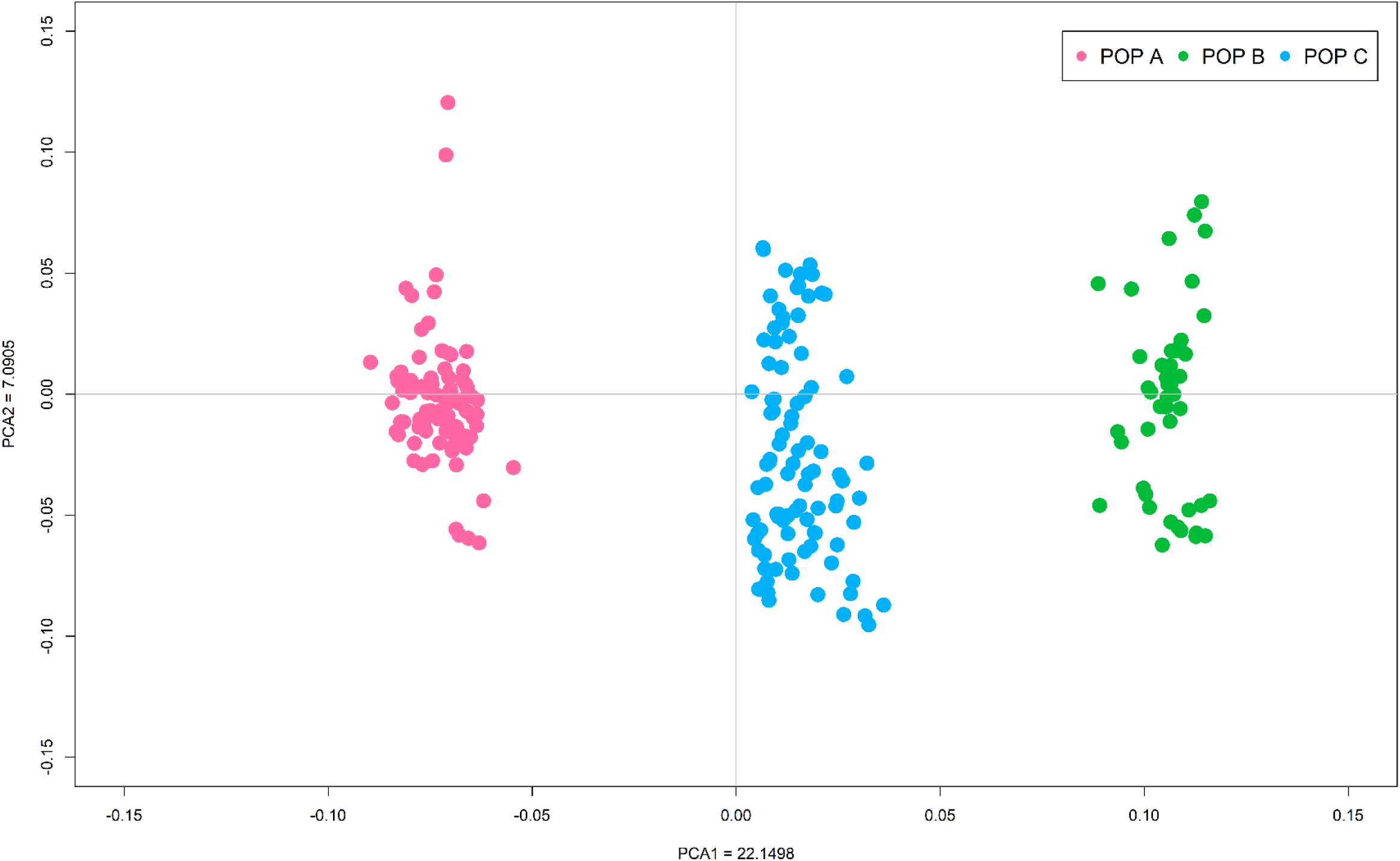
Principal component analysis of the autosomal genotypic data of three coho salmon populations.

The best K-value for admixture analysis was selected after performing several runs of MCMC for each K-value (ranging from 1 to 12), based on the posterior probability (Pr) of the fitted admixture model to the data with each K-value used (Pr(K)) [38]. The best K-value was suggested to be K = 11. These results indicate that POP A and B shared a large proportion of their genome with each other, probably due to the similar origin of the base populations. In addition, Figure 2 indicates a higher admixture level for POP C, due to the recent cross between POP A and B to generate this population.

**Figure 2.**
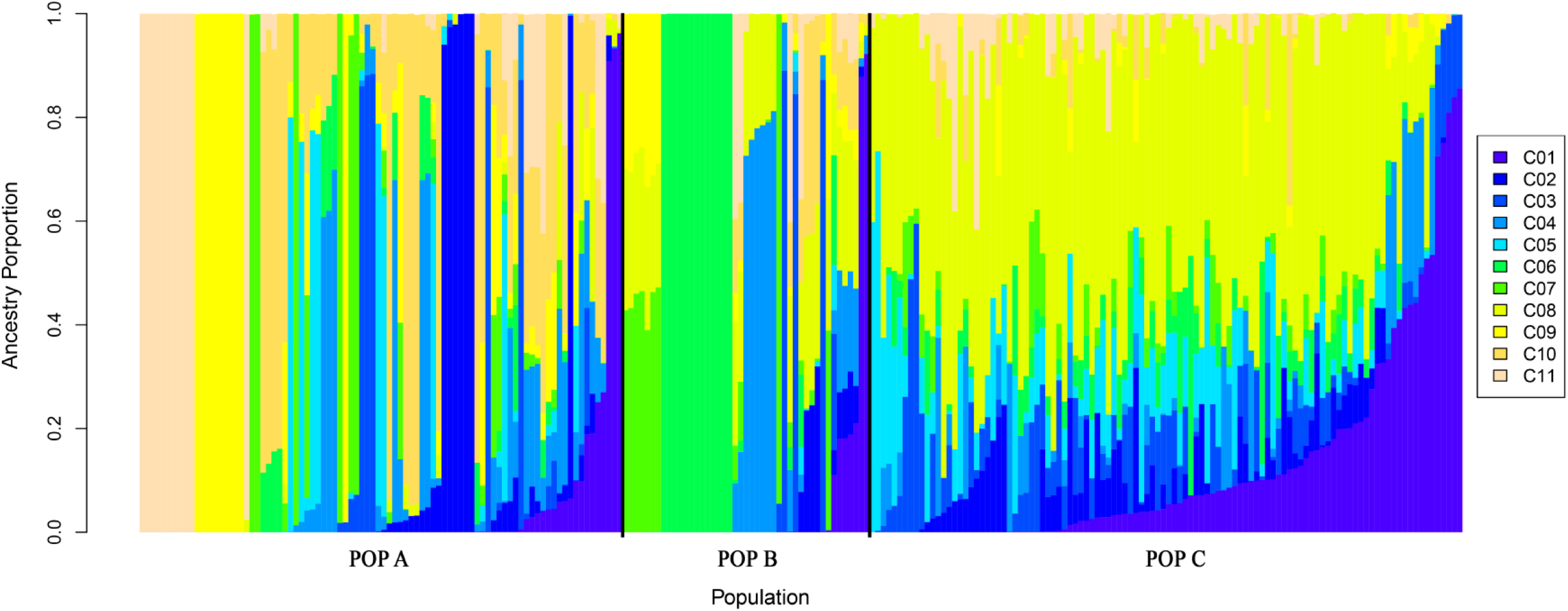
Admixture clustering of the three coho salmon population for K = 11.

### 3.2 Distribution of runs of homozygosity

We identified ROH in all animals for coho salmon POP A and B, and in 103 out of 108 individuals for POP C. A total of 3,250, 1,605, and 273 ROH and an average number of 36.93±7.13, 35.65±8.64, 2.65±1.27 ROH per animal were identified for POP A, B and C, respectively. The mean ROH length was 6.47±7.38, 7.172±7.69 and 2.58±2.07 Mb for POP A, B and C, respectively (Table 2) and the longest segment identified was 61.82 Mb, found in chromosome 2 for POP B (Figure S1). The ROH analysis for different length classes suggests that for the three coho salmon populations the genome is mostly composed of a high number of short segments (ROH_1–2 Mb_, ROH2–4 Mb). No segment was found for ROH>16Mb in POP C.

**Table 2.** Number of individuals (n), estimated of mean and standard deviation (SD) of inbreeding coefficient using runs of homozygosity (ROH) for different ROH length, based on excess of homozygosity (F_HOM_), genomic relationship matrix (F_GRM_) and pedigree-based relationship matrix (F_PED_),

| Class | POP A |  |  | POP B |  |  | POP C |  |  |
| --- | --- | --- | --- | --- | --- | --- | --- | --- | --- |
|  | n | Mean | SD | n | Mean | SD | n | Mean | SD |
| **ROH<sub>ALL</sub>** | 88 | 0.143 | 0.038 | 45 | 0.143 | 0.062 | 108 | 0.009 | 0.002 |
| **ROH<sub>1-2 Mb</sub>** | 88 | 0.120 | 0.143 | 45 | 0.143 | 0.061 | 102 | 0.009 | 0.022 |
| **ROH<sub>2-4 Mb</sub>** | 88 | 0.113 | 0.133 | 41 | 0.142 | 0.053 | 51 | 0.009 | 0.024 |
| **ROH<sub>4-8 Mb</sub>** | 88 | 0.100 | 0.115 | 43 | 0.126 | 0.050 | 42 | 0.011 | 0.026 |
| **ROH<sub>8-16 Mb</sub>** | 86 | 0.081 | 0.090 | 41 | 0.107 | 0.042 | 11 | 0.030 | 0.035 |
| **ROH<sub>>16 Mb</sub>** | 88 | 0.051 | 0.056 | 41 | 0.064 | 0.036 | 4 | 0.039 | 0.025 |
| **F<sub>HOM</sub>** | 88 | -0.036 | 0.048 | 41 | -0.058 | 0.082 | 108 | -0.106 | 0.028 |
| **F<sub>GRM</sub>** | 88 | 0.145<sup>b</sup> | 0.037 | 45 | 0.193<sup>a</sup> | 0.040 | 108 | 0.051<sup>c</sup> | 0.009 |
| **F<sub>PED</sub>** | 88 | 0.071 | 0.021 | 45 | 0.081 | 0.014 | 108 | 0.000 | 0.000 |

The number of ROH identified differs between chromosome and population. POP A has the highest number of ROH per chromosome, with more than 150 for chromosomes Okis5, Okis6 and Okis17. For POP B, chromosomes Okis5, Okis18 and Okis19 have more than 100 ROH, whereas for POP C, with the exception of chromosome Okis5, have less than 50 per chromosomes (Figure 3). The average ROH length also differs between chromosomes and population. POP A has two chromosomes (Okis5 and Okis11) with ROH segments greater than 10 Mb. POP B has five chromosomes (Okis3, Okis4, Okis6, Okis11 and Okis14) with ROH segments greater than 10 Mb; while all chromosomes in POP C have ROH segments smaller than 7 Mb (Figure 4).

**Figure 3.**
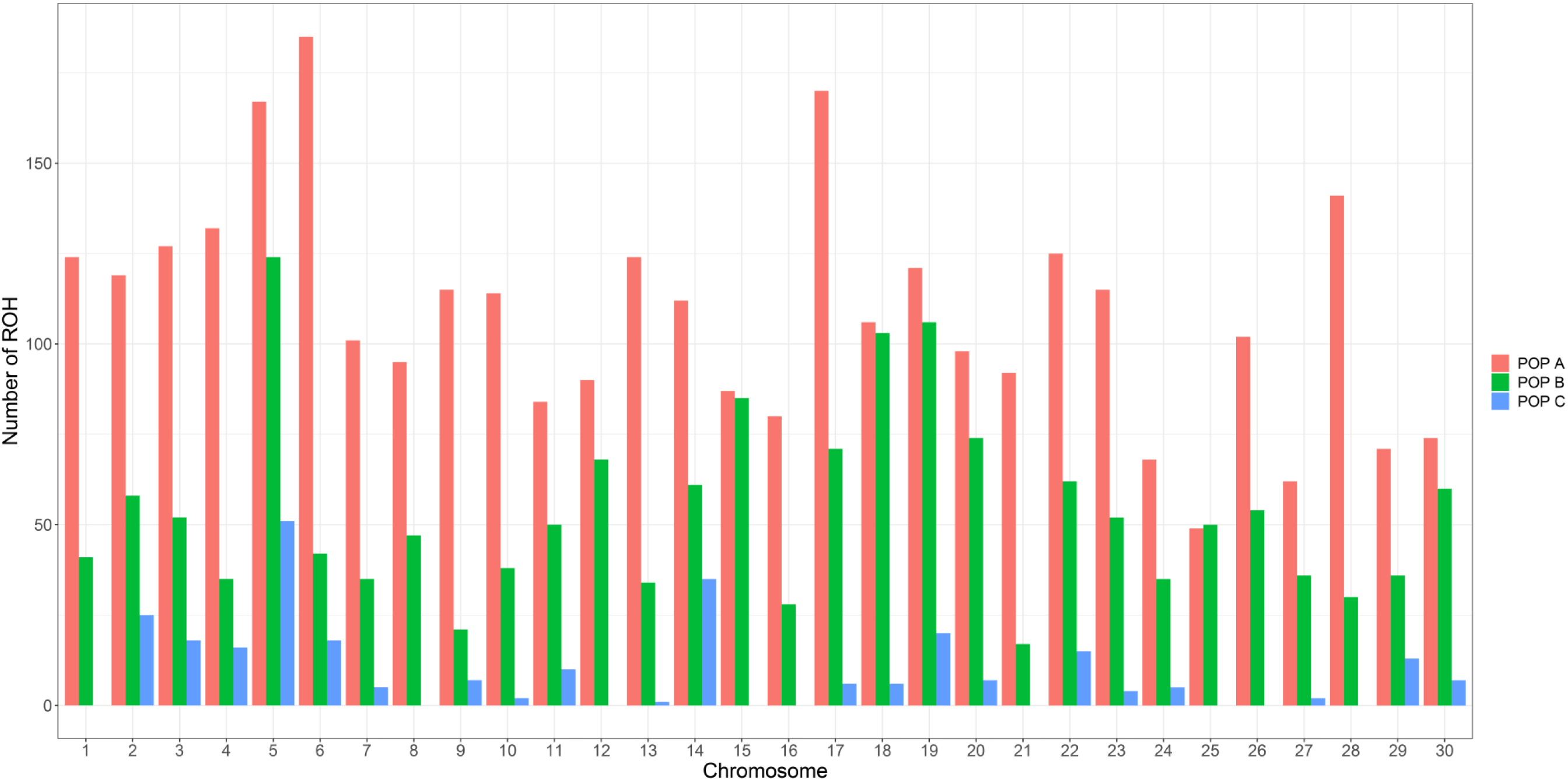
Distribution of number of runs of homozygosity (ROH) for each chromosome in three coho salmon populations.

**Figure 4.**
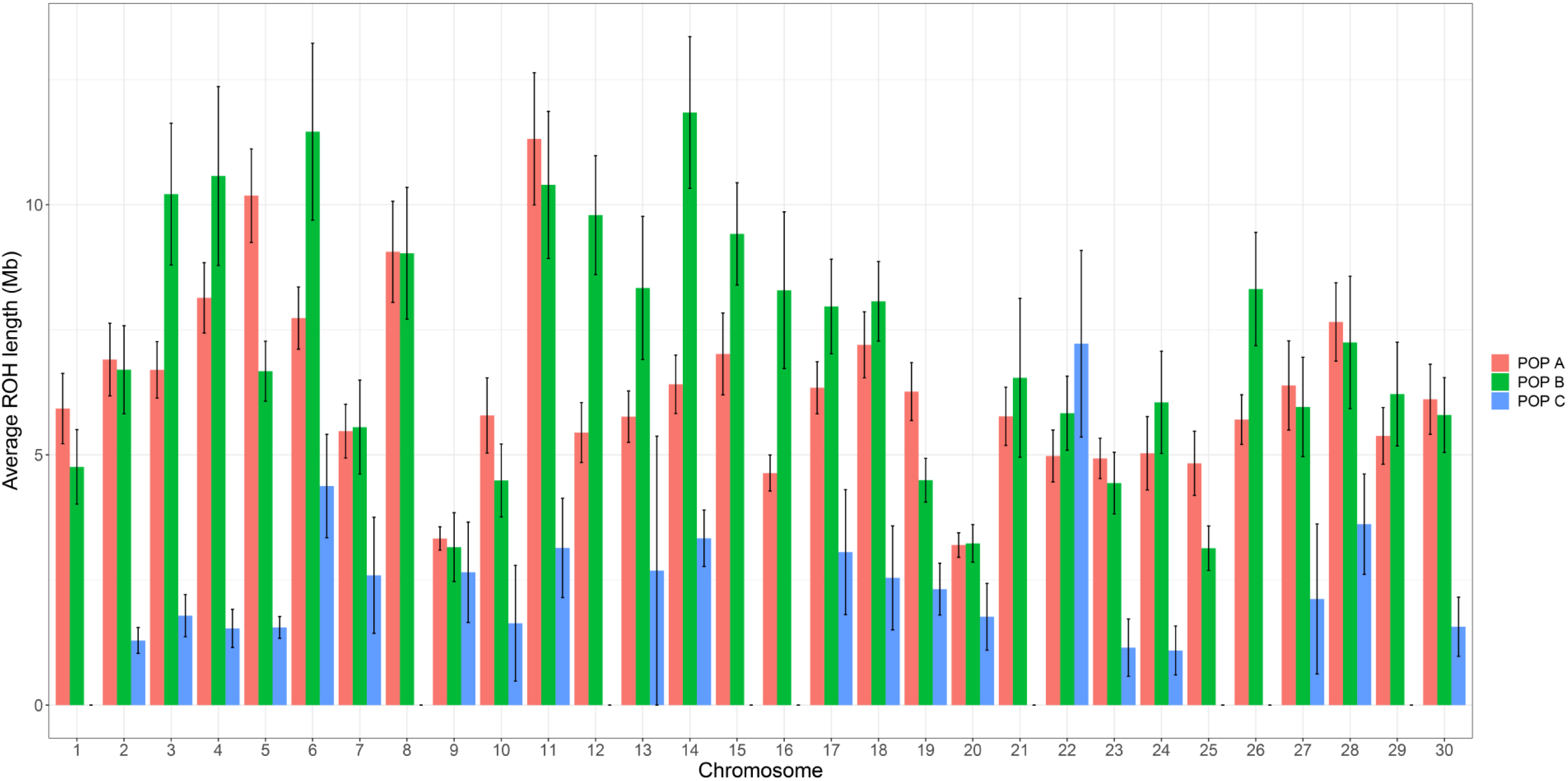
Average runs of homozygosity (ROH) length and standard error bars for each chromosome in three coho salmon populations.

Figure 5 shows the relation between the total number of ROH and the total length of ROH for each animal across the three populations. A considerable difference between POP C and POP A or B was found. For POP C, all animals have a small number of ROH (<8) with total length <25 Mb, whereas most individuals in POP A and B have at least 20 ROHs with a total length >100 Mb, with some extreme individuals with segments covering more than 300 Mb. The number of ROHs and segment length per animal and per chromosome are shown in Figure S1. The high number of segments >10 Mb in Okis5, Okis6 and Okis28, especially for POP A and B, suggests recent events of inbreeding, whereas the small segments as in Okis20 for POP A and B, and for most of chromosomes for POP C, suggests more ancient inbreeding.

**Figure 5.**
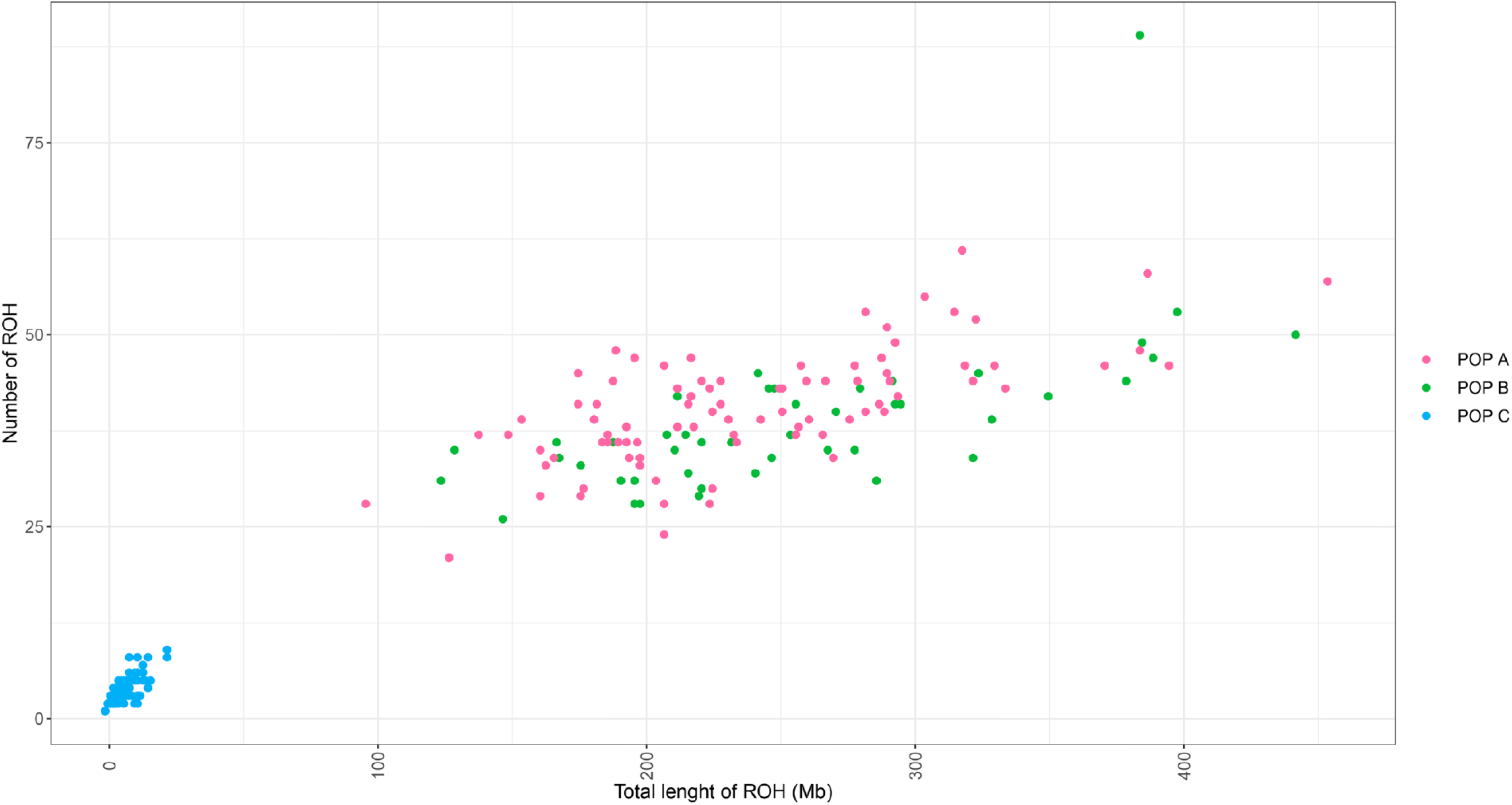
Relationship between the number of runs of homozygosity (ROH) and total length of ROH (Mb) per individual from each population.

### 3.3 Genomic- and pedigree-based inbreeding

We used four different methods to estimate the inbreeding coefficient, from the information of 102K markers and pedigree data (Table 3). The average inbreeding coefficient estimated using ROH was different between ROH classes, the values decreased when the ROH length segments increased for all populations. The mean value for FROH_ALL_ was the same for both POP A and B (0.142 and 0.152, respectively), but it was significantly different (p<0.05) for POP C (0.004) when compared to POP A or B. The F_HOM_ resulted in the lowest inbreeding values ranging from −0.036 to −0.105 for POP A and C, respectively. The mean value for F_GRM_ was different (p<0.05) between the three populations, the highest and lowest values were reported for POP B and C, respectively, whereas the F_PED_ value was not different between POP A and B, but was significantly lower for POP C (0.002, p<0.05). Additionally, we calculated the inbreeding coefficient based on the ROH per chromosome (Figure 6). POP A and B had the most chromosomes with inbreeding values higher than 0.2, as in Okis5, Okis6 and Okis28 for POP A, and Okis5, Okis12, Okis14, Okis18 and Okis26 for POP B, whereas lower values were found for POP C and for most of the chromosomes the inbreeding was equal to zero.

**Figure 6.**
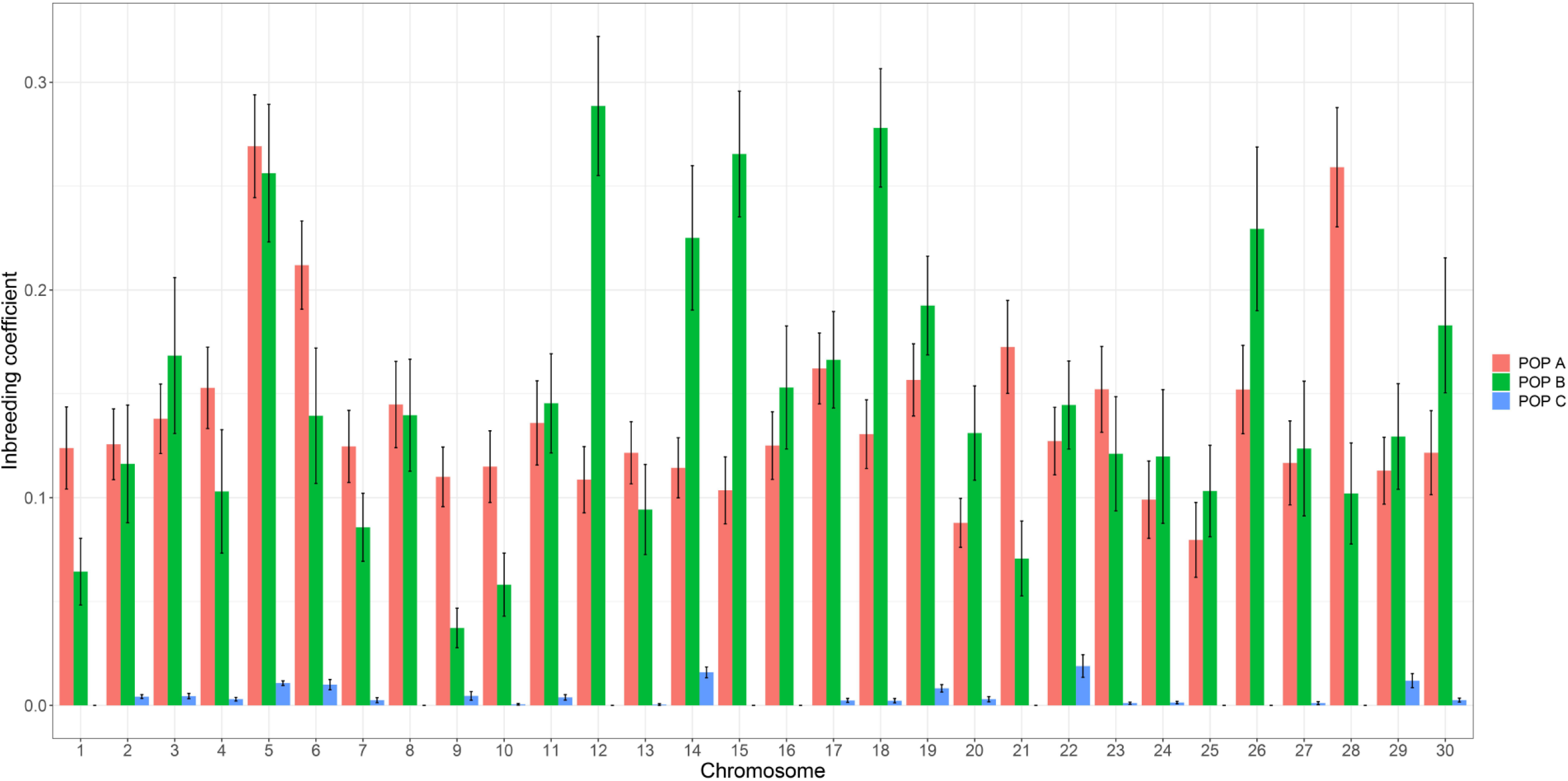
Distribution of inbreeding coefficients estimated using runs of homozygosity (ROH) for each chromosome in three coho salmon populations. Standard error bars were computed among individuals from the same population.

The Pearson correlation between different genomic methods to estimate the inbreeding coefficient suggested a high positive correlation (>0.82, p<0.001) for POP A and POP B (Figure 7 and 8, respectively). Correlation between different ROH length classes decreased in function with the comparison between shorter and longer segments, e.g. highest correlation between ROH_1–2 Mb_ and ROH_2–4 Mb_ and lowest between ROH_1–2 Mb_ and ROH_>16Mb_. The lowest correlation values among genomic methods was reported between ROH_>16Mb_ and both ROH_HOM_ and ROH_GRM_. In addition, for POP A and POP B correlation low correlation values were found, respectively, ranging from 0.35 to 0.39 (p<0.01), between genomic methods and F_PED_.

**Figure 7.**
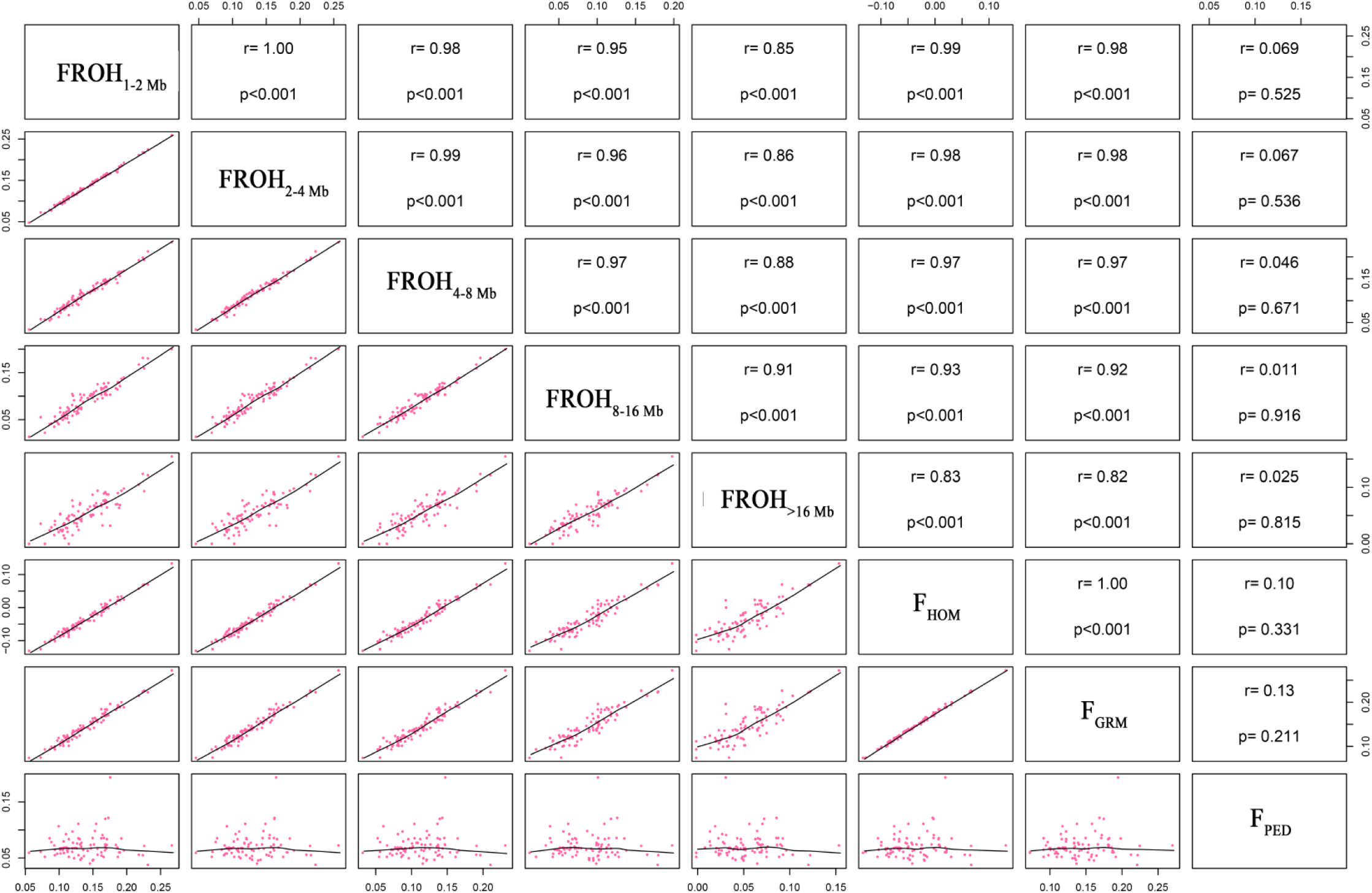
Scatterplots (lower panel) and Pearson correlations (upper panel) of genomic inbreeding coefficients using runs of homozygosity (ROH) for different ROH length, based on excess of homozygosity (F_HOM_), genomic relationship matrix (F_GRM_) and pedigree-based relationship matrix (F_PED_) for POP A.

**Figure 8.**
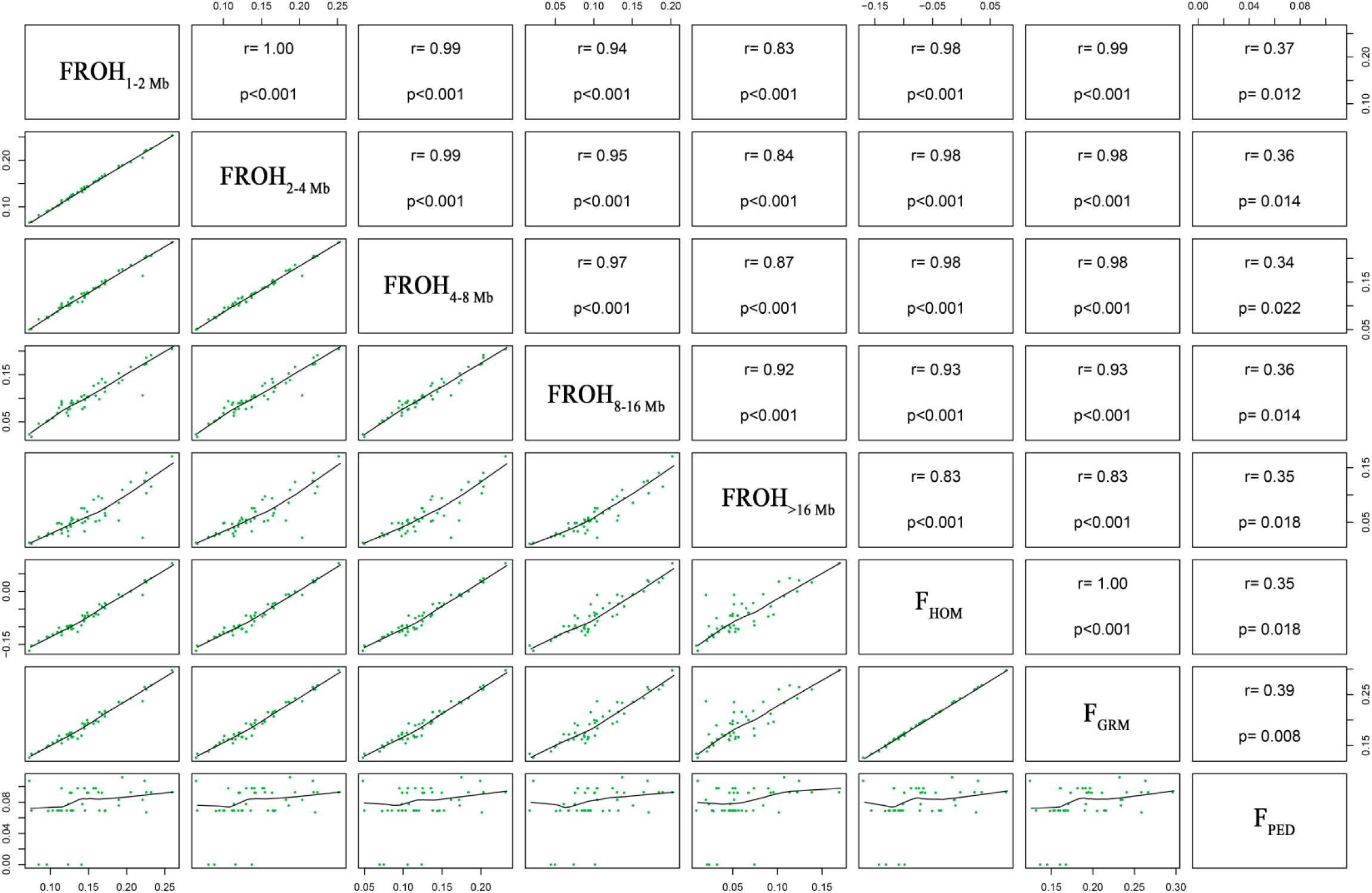
Scatterplots (lower panel) and Pearson correlations (upper panel) of genomic inbreeding coefficients using runs of homozygosity (ROH) for different ROH length, based on excess of homozygosity (F_HOM_), genomic relationship matrix (F_GRM_) and pedigree-based relationship matrix (F_PED_) for POP B.

Different patterns of correlations were observed for POP C, compared to POP A and B, probably due to the low inbreeding level of this recently admixed population. Medium to high positive correlation was reported between the ROH classes (0.54 to 0.94, p<0.001), and a correlation equal to unity was observed between ROH_HOM_ and ROH_GRM_. For other correlations, small values (ranged from 0.28 to 0.34) or not different from zero were observed (Figure 9).

**Figure 9.**
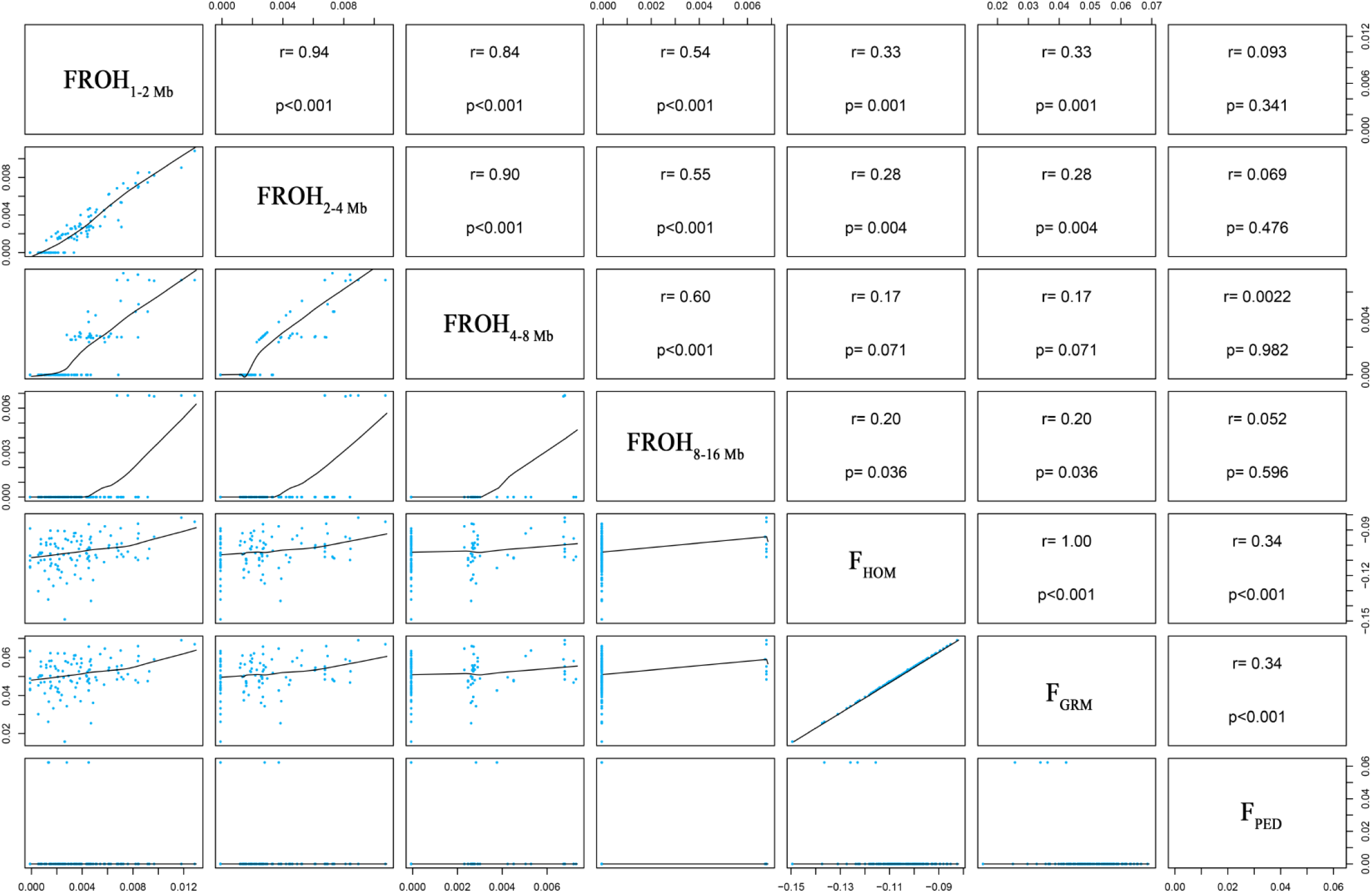
Scatterplots (lower panel) and Pearson correlations (upper panel) of genomic inbreeding coefficients using runs of homozygosity (ROH) for different ROH length, based on excess of homozygosity (F_HOM_), genomic relationship matrix (F_GRM_) and pedigree-based relationship matrix (F_PED_) for POP C.

## 4. Discussion

### 4.1 Genomic population structure

The first two principal components explained more than 29% of the total genetic variation for the three populations studied, which were separated into three different clusters (Figure 1). The admixture results are in agreement with the recent event of hybridization of POP A and B to generate POP C, where the genetic differentiation between POP A and B may have been be partly generated by differences in the base population, which can have a pronounced effect on allele frequencies [42]. In addition, considering that POP A and B have been independently selected by at least eight generations each, differences in the selection processes, as well as the environmental conditions and drift, may have influenced the differences observed in Figure 2.

### 4.2 Runs of homozygosity characterization

Figure 3 and 4 show that independent of the population, the ROH patterns seem to be differentially distributed within specific genomic regions, same as the inbreeding values between chromosomes (Figure 6). The highest autozygosity, e.g. in chromosome Okis5 and Okis6 for POP A and B, is likely the consequence of artificial selection [26], considering that these populations have been under genetic selection for harvest weight for at least eight generations. A ROH study in humans [43] suggested that the homozygosity segments are more common in regions with high linkage disequilibrium (LD) and low recombination rates. Thus the highest mean levels of LD found in Okis5 and Okis6 in animals from the same populations [35] are in accordance with the two chromosomes with the highest number of ROH in the present study.

Differences in the number of ROH and segment length was observed within and across populations (Figure 5 and Additional file 1). The higher number of ROH in POP A compared to POP B is most likely due to higher sample size in the former, whereas the differences in ROH length between the three populations may be due to differences in the effective population size, selection intensities, or threshold of inbreeding allowed for the matings, suggesting that artificial selection commonly increases the autozygosity across the genome and creates long ROH in specific regions of the genomes [26]. In contrast, the shorter segments and smaller number of ROH in POP C when compared against both POP A and B may be the result of recent population admixture between these populations. Furthermore, animals from the same population might have the same total ROH lengths but a variable number of segments, which is probably the result of different distances from common ancestors [25]. Interestingly, for both POP A and B, the length class ROH_2-4 Mb_ has more ROH than ROH_1-2 Mb_ (Table 2), which is different than what is commonly found in other species [15,44,45]. These differences can be due to the criteria adopted to identify ROH or an inherent characteristic of these populations. There is no consensus on the best parameters to characterize ROH patterns [32]; thus, here we used the minimum number of 50 SNPs and the length of 1 Mb to define a ROH segment. We chose the current parameters due to the historical demographics of coho salmon in Chile. The ROH_2-4 Mb_ should date from about 20 generations ago (approximately 40 years considering the generation interval of 2 years), which corresponds to the introduction of coho salmon in Chile at the end of the 1970s, to begin the establishment of Chilean brood stocks [2,35].

### 4.3 Genomics- and pedigree-based inbreeding

Based on information of ROH length it is possible to infer the number of generations for inbreeding events [46]. The ROH due to ancient origin tend to be shorter, e.g. ROH_1–2 Mb_, ROH_2–4 Mb_ and ROH_4-8 Mb_ date from 50, 20 and 12.5 generations ago, respectively. In contrast, recent ROH are longer, due to the small probability of breaking down the segments that are identical-by-descent (IBD) by means of recombination events. Thus, the ROH_8–16 Mb_ and ROH_>16 Mb_ are dated to 6 and 3 generations ago, respectively [22,46]. For both POP A and B it was possible to identify short and long segments in most of the animals analyzed, whereas in the POP C a small number of animals (n = 7) presented ROH_8–16 Mb_ and none ROH_>16 Mb_.

In recent years, some studies have investigated different genomic methods to estimate inbreeding coefficients in cattle [12,25,26,45,47,48], pigs [27,28,49,50], goats [51–53] and rainbow trout [32]. However, this is the first study aimed at characterizing the ROH patterns and comparing different genomic- and pedigree-based methods to estimate inbreeding coefficients in farmed coho salmon populations. Both genomic- and pedigree-based strategies have some advantages and disadvantages. The pedigree inbreeding coefficient, is a simple method that requires recording genealogy information, but does not account for the autozygosity differences among animals with the same inbreeding history. In contrast, genomic inbreeding can measure the realized inbreeding of an individual and incorporate the breeding history of the animal, including new mutations, ancient and contemporary inbreeding [27].

A comparison of inbreeding coefficients, showed F_GRM_ gave the highest values, especially for B and C, probably because the alleles IBD and identical by state (IBS) are not differentiated for F_GRM_ [12]. This result is in agreement with results previously found in humans, cattle, and simulation studies [12,15,16]. F_HOM_ resulted in negative inbreeding values for all populations (Table 3), suggesting that the individuals have lower levels of homozygosity than expected in the reference population under Hardy-Weinberg equilibrium [54] and underestimated values should be expected [55]. The F_PED_ for POP A and POP B were smaller than values estimated using FROH_ALL_ and F_GRM_, but are in accordance with the values estimated for the same populations using previous generations [8,10]. The F_PED_ can be easily underestimated when pedigree information of less than 20 generations is used [55]. The difference between F_ROH_ and F_PED_ could be also due to the unknown pedigree information before recording, which in practical terms means that inbreeding levels for founding animals were not zero.

### 4.4 Inbreeding coefficients correlations

ROH can be identified for each animal, and the inbreeding coefficient will reflect the direct level of homozygosity, not influenced by allele frequencies [19]. Also with regard to information about recent and remote inbreeding [56]. High correlations (>0.80) were found between F_ROH_ and other genomic inbreeding estimates for POP A and B (Figure 7 and 8). Some authors have also reported a strong or moderate correlation between genomics methods used to calculate inbreeding coefficients for different species [11,27,57,58].

The genomic-based inbreeding method correlated moderately or poorly with pedigree data, showing values lower than 0.39 (Figure7 to 9). Similary weak or no correlation was reported for cattle [24,45,47], whereas a moderate to strong positive correlation was described by some authors [15,40,48,59]. An increase in the correlation between genomic- and pedigree-based inbreeding as the pedigree depth increases is expected [24]. Here we used the complete pedigree information of nine generations for both POP A and POP B, whereas for POC C a pedigree depth of eight generations was used. In a previous pedigree-based inbreeding study using the same broodstock population of POP A (7^th^ generation) and POP B (8^th^ generation), an increasing tendency for inbreeding values in the last four generations was reported for both populations [8] and a continued inbreeding accumulation until 9^th^ generation used in our study is well-known. Thus, we expected a higher correlation between long ROH segments (ROH_8–16 Mb_, and ROH_>16 Mb_) and F_PED_ values. The weak or no correlation may be explained by the depth of pedigree records [55], incorrect or incomplete pedigree information [47], the F_PED_ that assumed the founder individuals are unrelated [12], and the fact that F_PED_ does not consider the stochastic nature of recombination and the persistence of ancestral short segments through time, due to the lack of recombination in specific regions [11]. These facts suggest that the F_PED_ may not reflect true inbreeding values. Additionally, the population sample size must be representative to avoid population stratification [15,24] and to improve the correlation between genomic- and pedigree-based inbreeding. However, in our case, POP B is the population with the smallest sample size (n = 45), but was the only one that resulted in significant correlations (Figure 8). Various studies of ROH used similar or smaller sample size in livestock species and rainbow-trout [19,32,44,48].

A relatively large effective population size (Ne) is recommended to maintain the control of inbreeding in the medium-term. However, decline in the historical Ne was reported for animals from the same population as POP A [35]. The reduction may be due to the prioritization of genetic gain using high selection pressure without putting strong control on the family contribution for each generation [8]. Consequently, mating close relatives is more probable, which results in a high level of inbreeding and the creation of long ROH segments for both POP A and B. Therefore, to increase the effective population size and to limit the inbreeding level [8], POP C was generated. According to our results, this strategy was effective in reducing the inbreeding levels and changing the patterns of ROH, clearly differentiating from POP A and B. These results are in accordance with some studies [15,48,60,61] that suggest that high heterogeneity populations due admixture or crossbreeding lines contributed to the breakdown of long homozygous segments and reduced the inbreeding levels in captive populations.

## 5. Conclusion

In this study, we found different numbers and lengths of runs of homozygosity in three coho salmon populations farmed in Chile. Moreover, the inbreeding coefficient estimated using genomic- or pedigree-based methods have varied among populations and the high correlations between genomic inbreeding methods suggest that these are the more accurate methods to estimate autozygosity levels and thus must be used as an alternative when pedigree information is inaccurate, incomplete or unavailable.

## Acknowledgements

We thank the government of Canada through Genome Canada, Genome British Columbia, and Genome Quebec to supported this research that was carried out in conjunction with the projected EPIC4 (Enhanced Production in Coho: Culture, Community, Catch).

## Authors’ contributions

GMY performed the analysis and wrote the initial version of the manuscript. PC and RMN contribute with writing. BK develop the chip array. JMY conceived and designed the study; contributed to the discussion and writing. All authors have reviewed and approved the manuscript.

## Competing interests

The authors declare that the research was conducted in the absence of any commercial or financial relationships that could be construed as a potential conflict of interest.

---

Figure S1. Runs of homozygosity patterns for all chromosome (Okis1 to Okis30) in three coho salmon population. Each row represents one individual and each bar a ROH segment.

