## Supplementary Figure 1 for "Estimates of autozygosity through runs of homozygosity in farmed coho salmon"

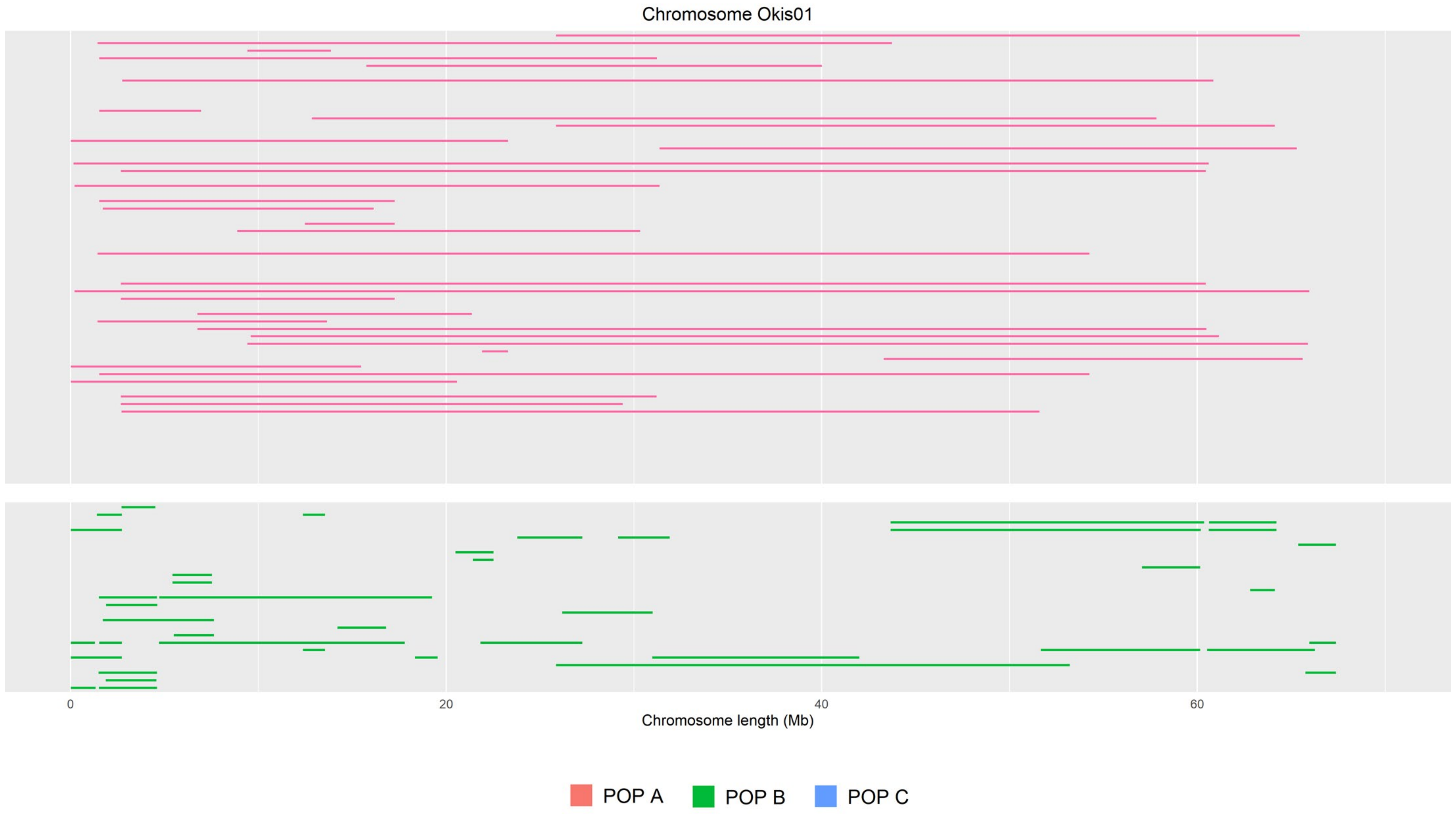
Chromosome Okis01
0
20
40
60
Chromosome length (Mb)
POP A
POP B
POP C

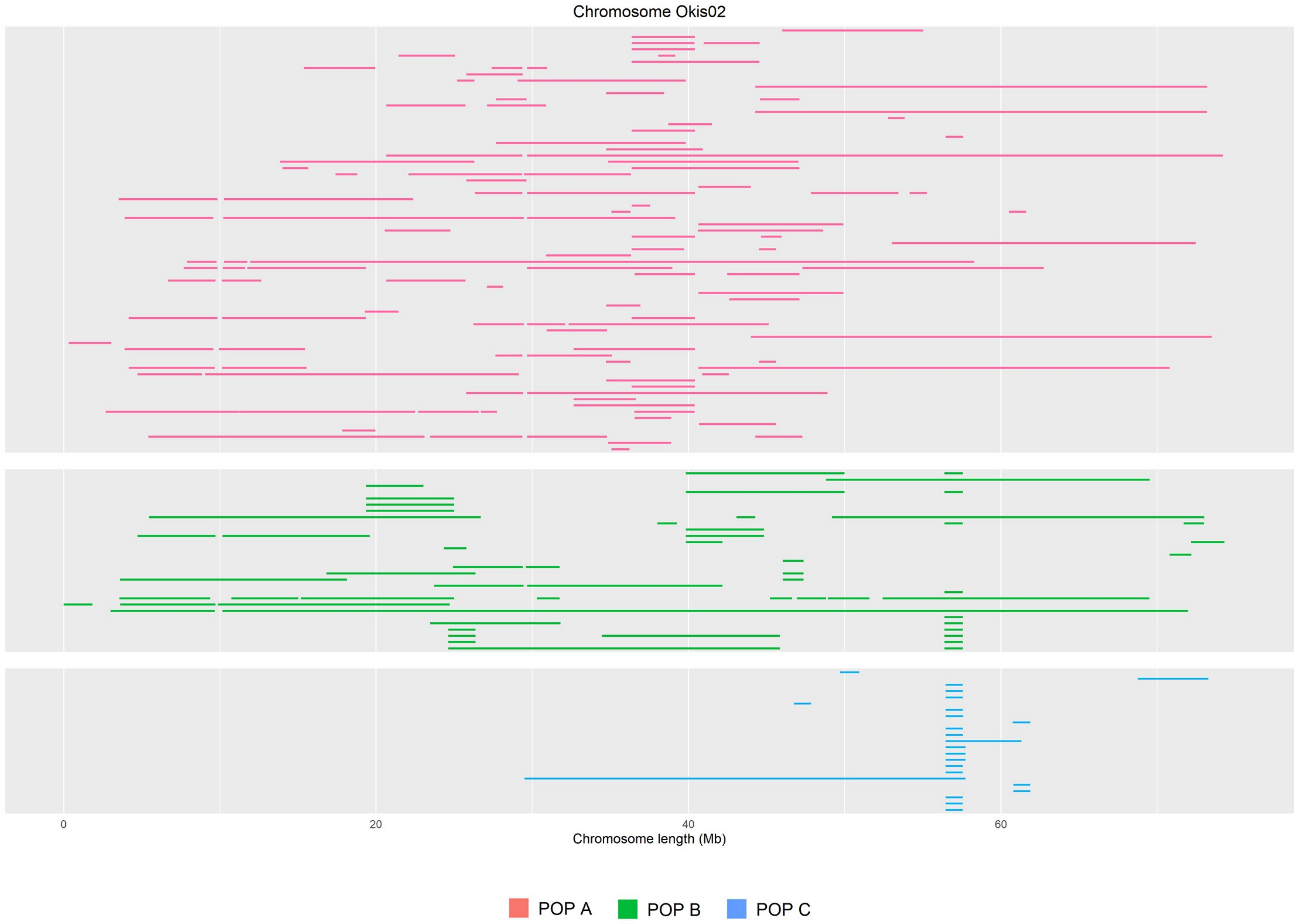

Chromosome Okis02
0
20
40
60
Chromosome length (Mb)
POP A
POP B
POP C

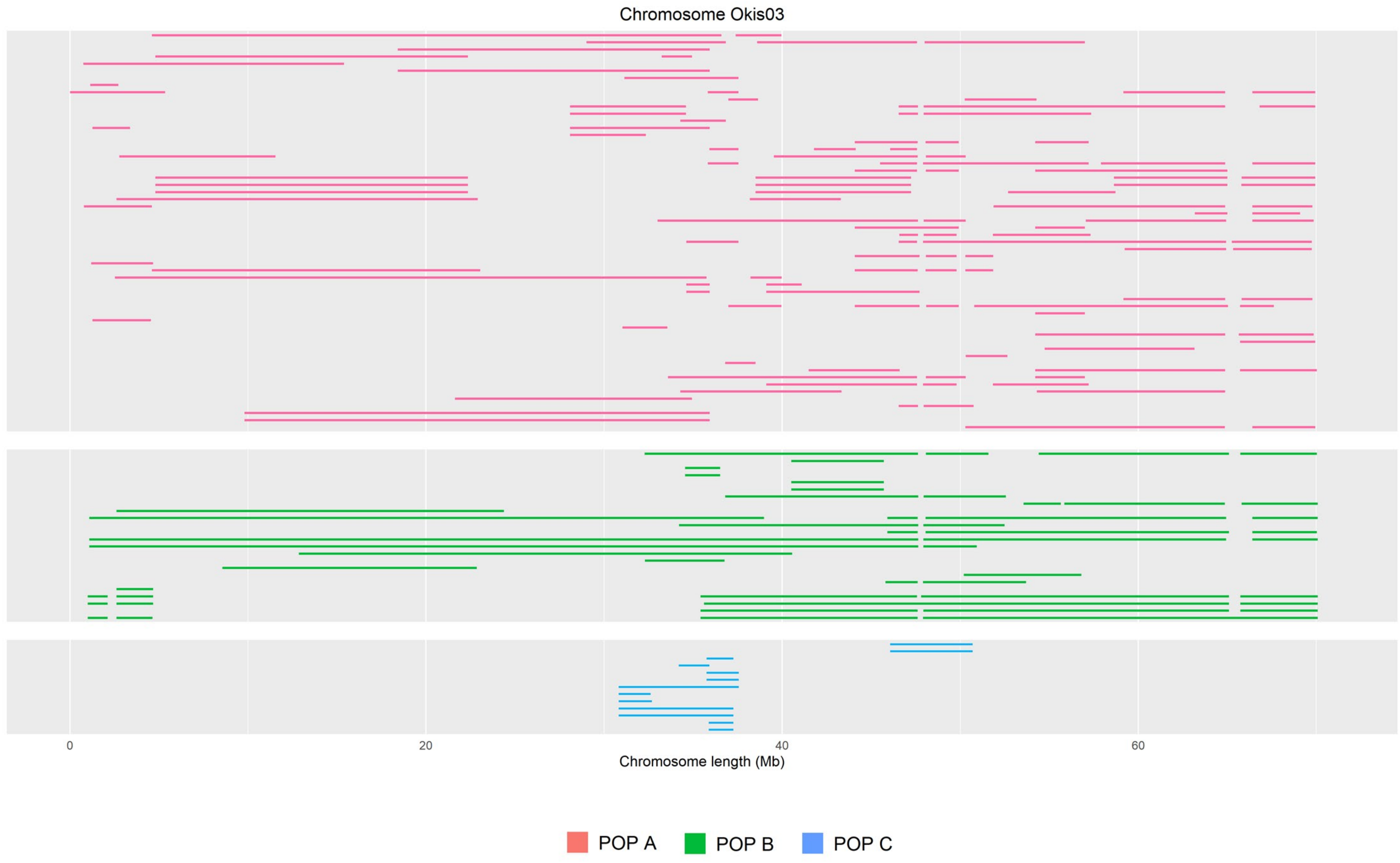
Chromosome Okis03
0
20
40
60
Chromosome length (Mb)
POP A
POP B
POP C

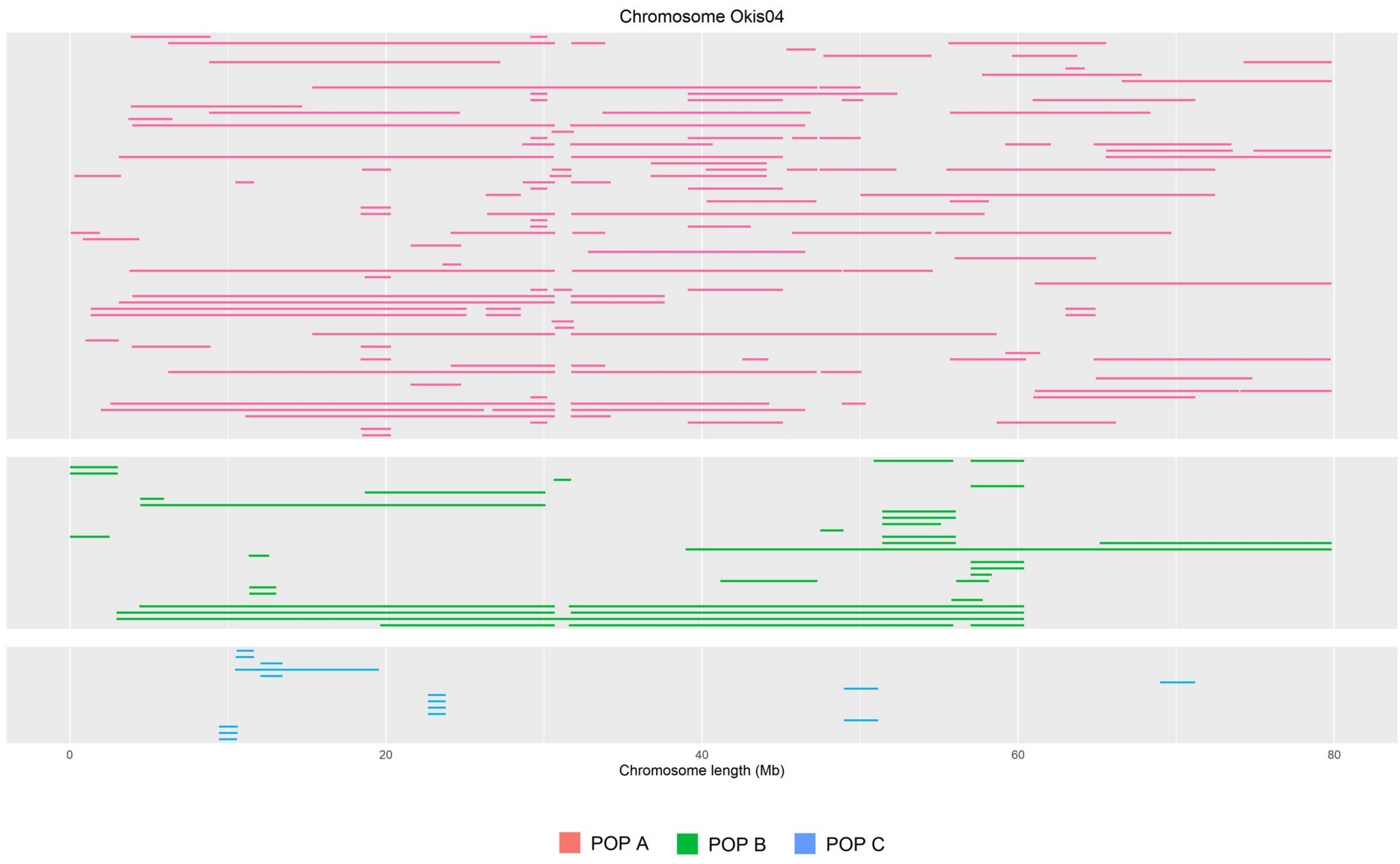
Chromosome Okis04
0
20
40
60
80
Chromosome length (Mb)
POP A
POP B
POP C

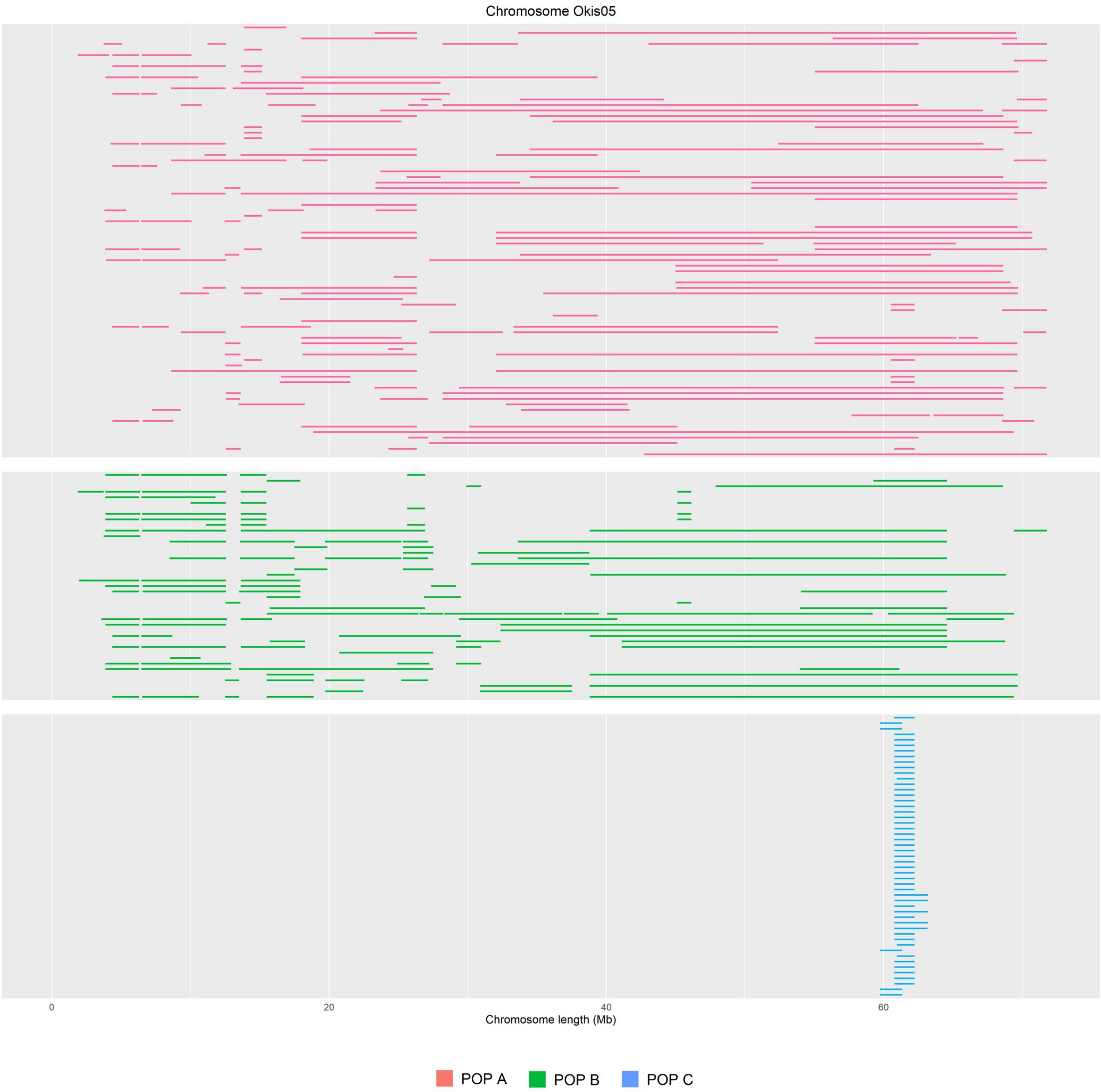
Chromosome Okis05
0
20
40
60
Chromosome length (Mb)
POP A
POP B
POP C

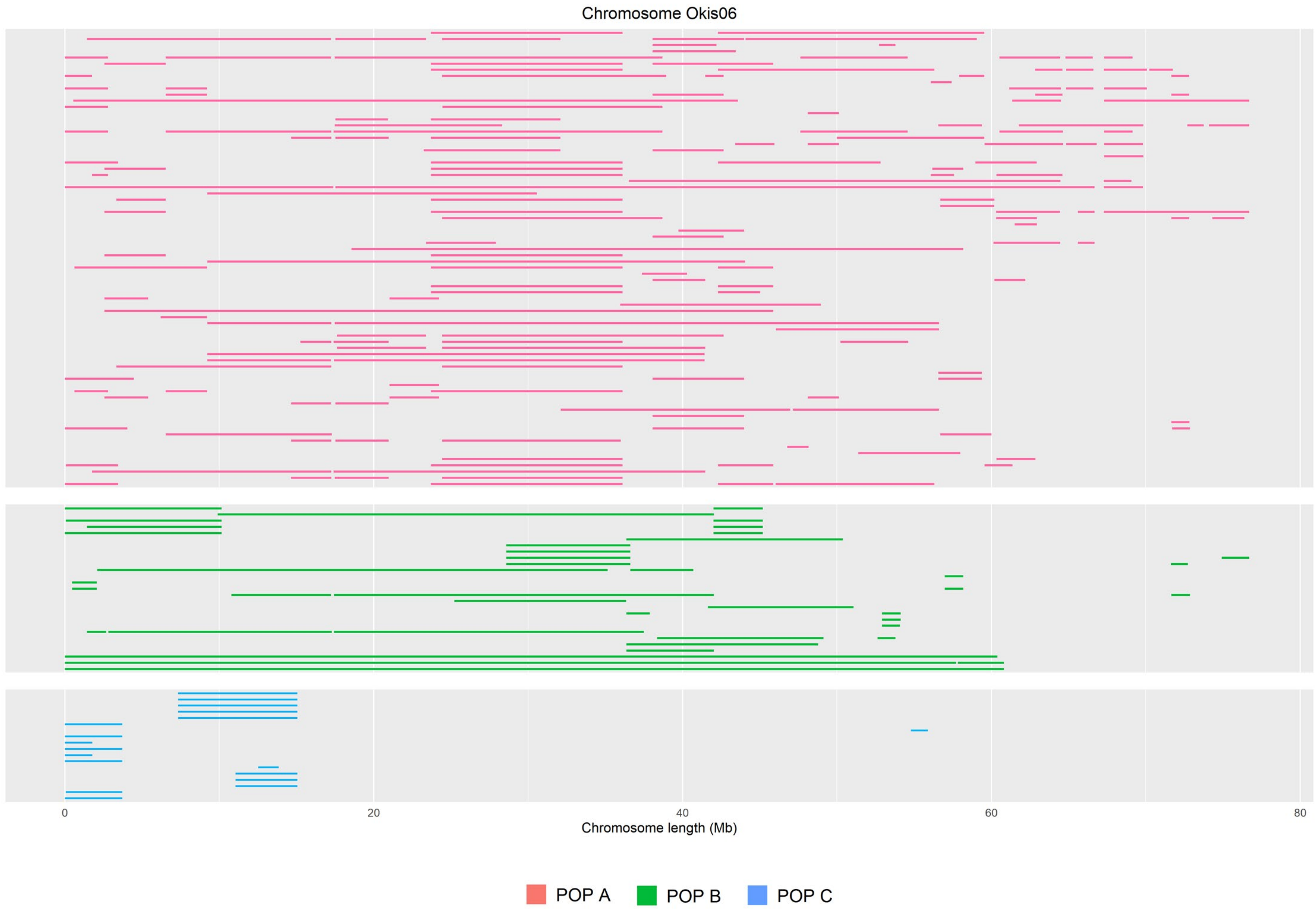
Chromosome Okis06
0
20
40
60
80
Chromosome length (Mb)
POP A
POP B
POP C

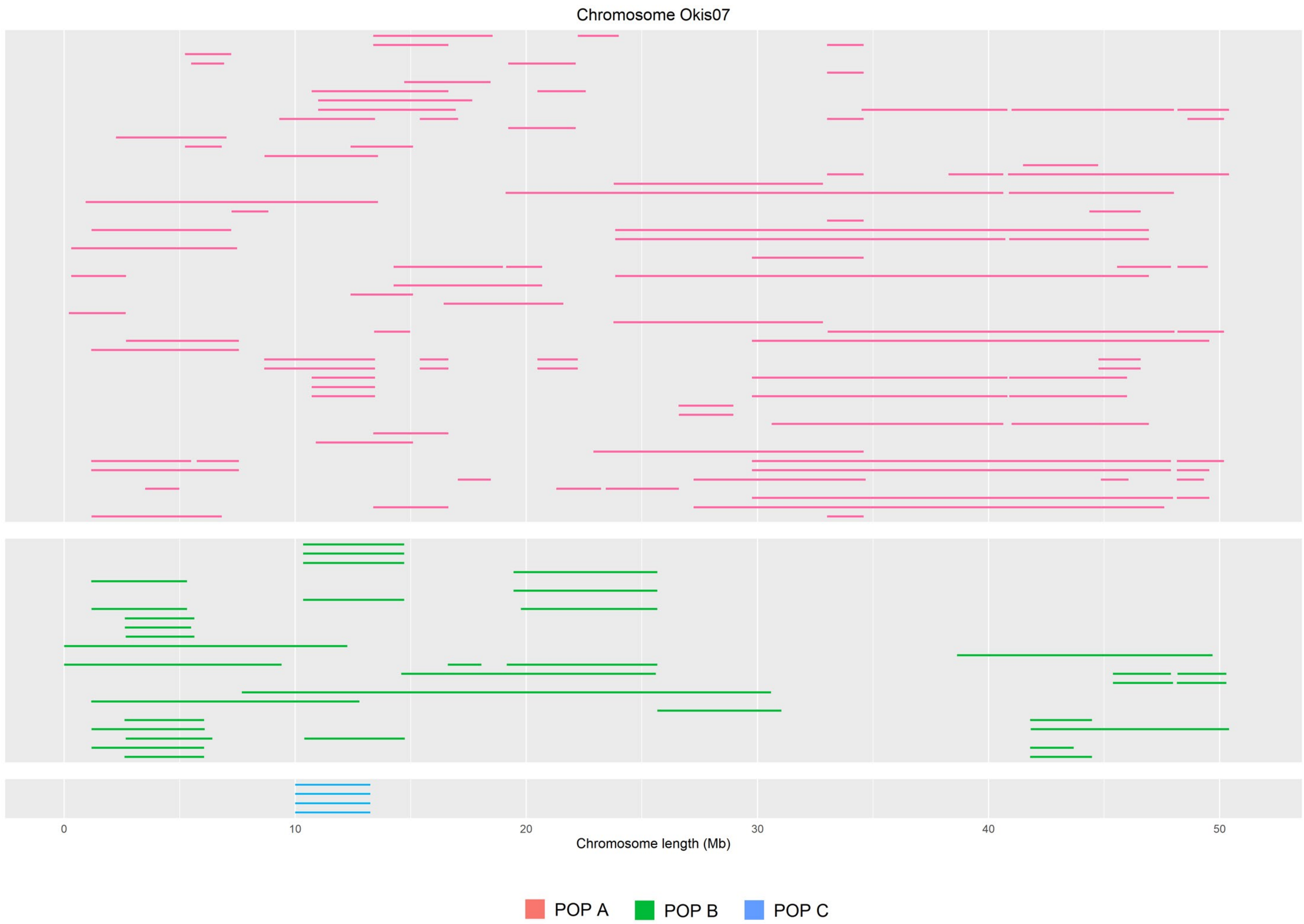

Chromosome Okis07
0
10
20
30
40
50
Chromosome length (Mb)
POP A
POP B
POP C

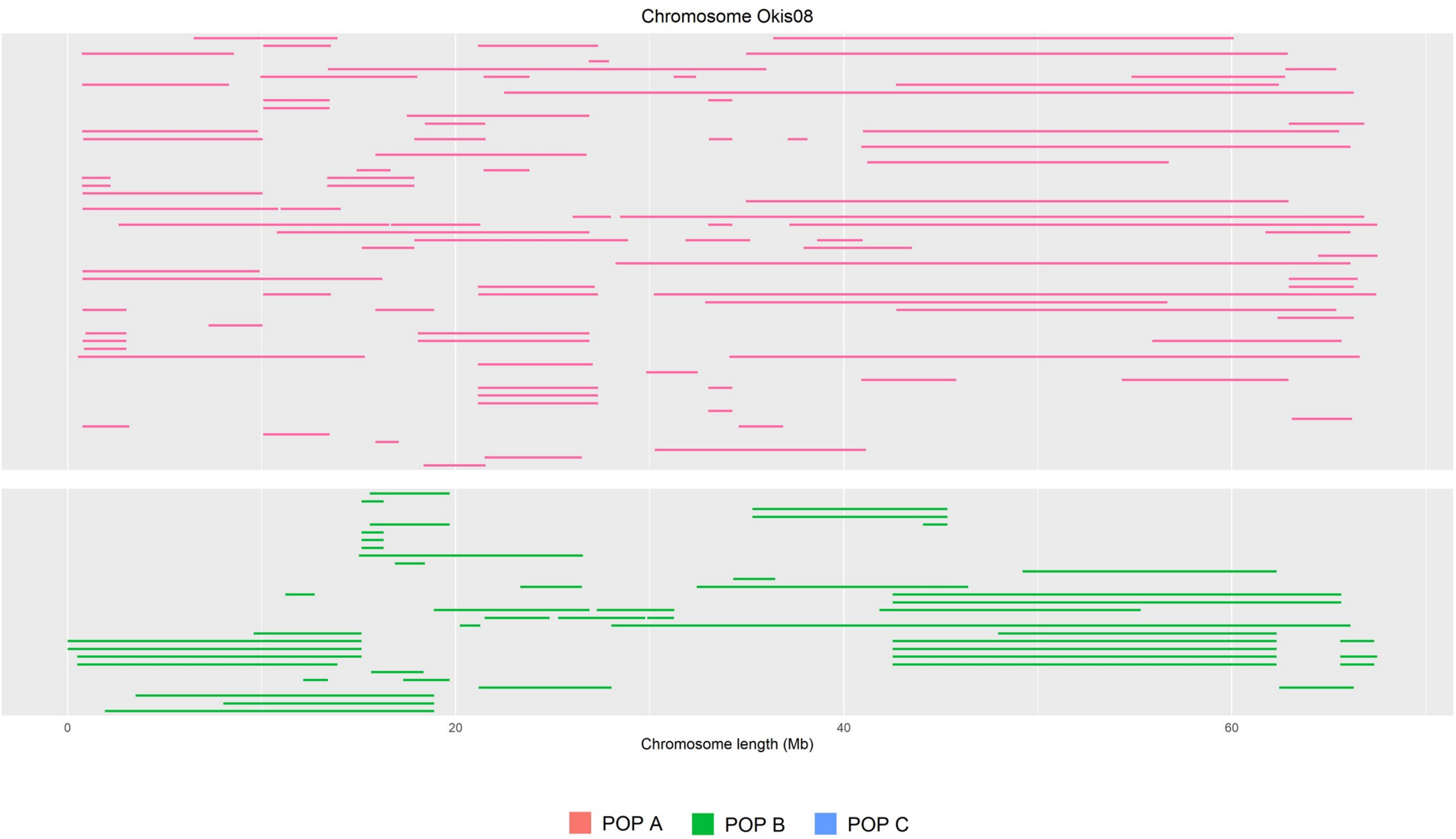
Chromosome Okis08
0
20
40
60
Chromosome length (Mb)
POP A
POP B
POP C

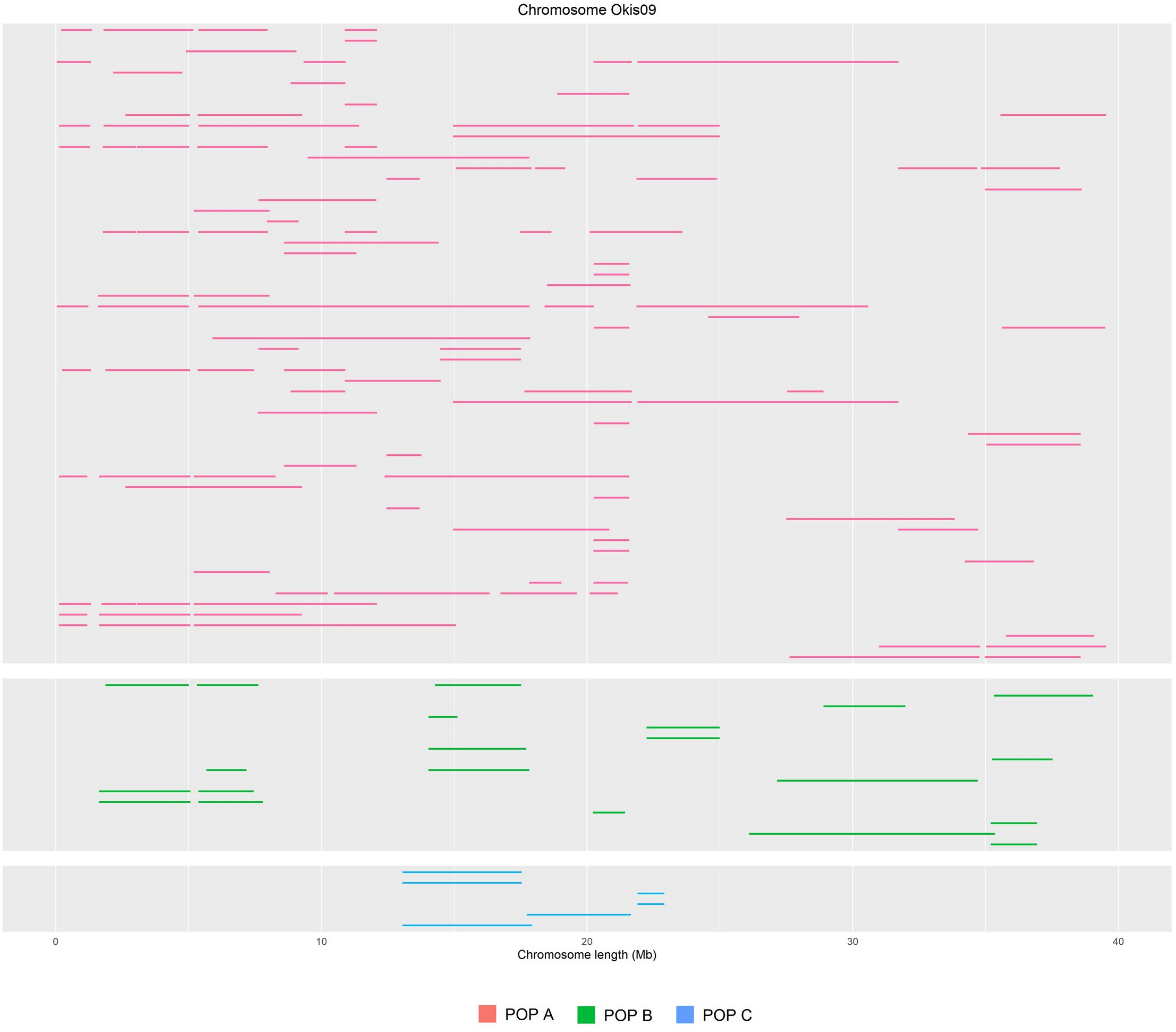
Chromosome Okis09
0
10
20
30
40
Chromosome length (Mb)
POP A
POP B
POP C

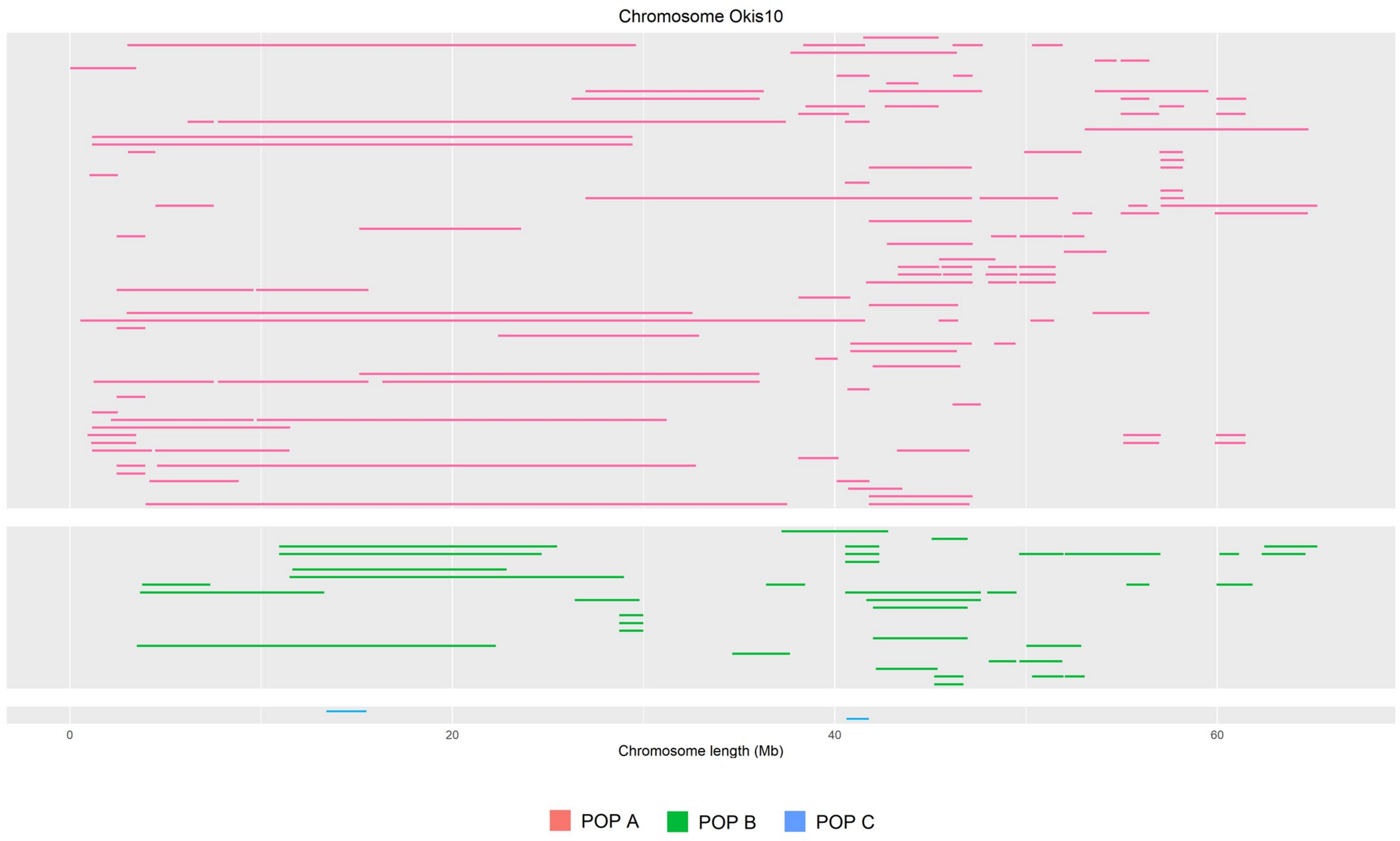

Chromosome Okis10
0
20
40
60
Chromosome length (Mb)
POP A
POP B
POP C

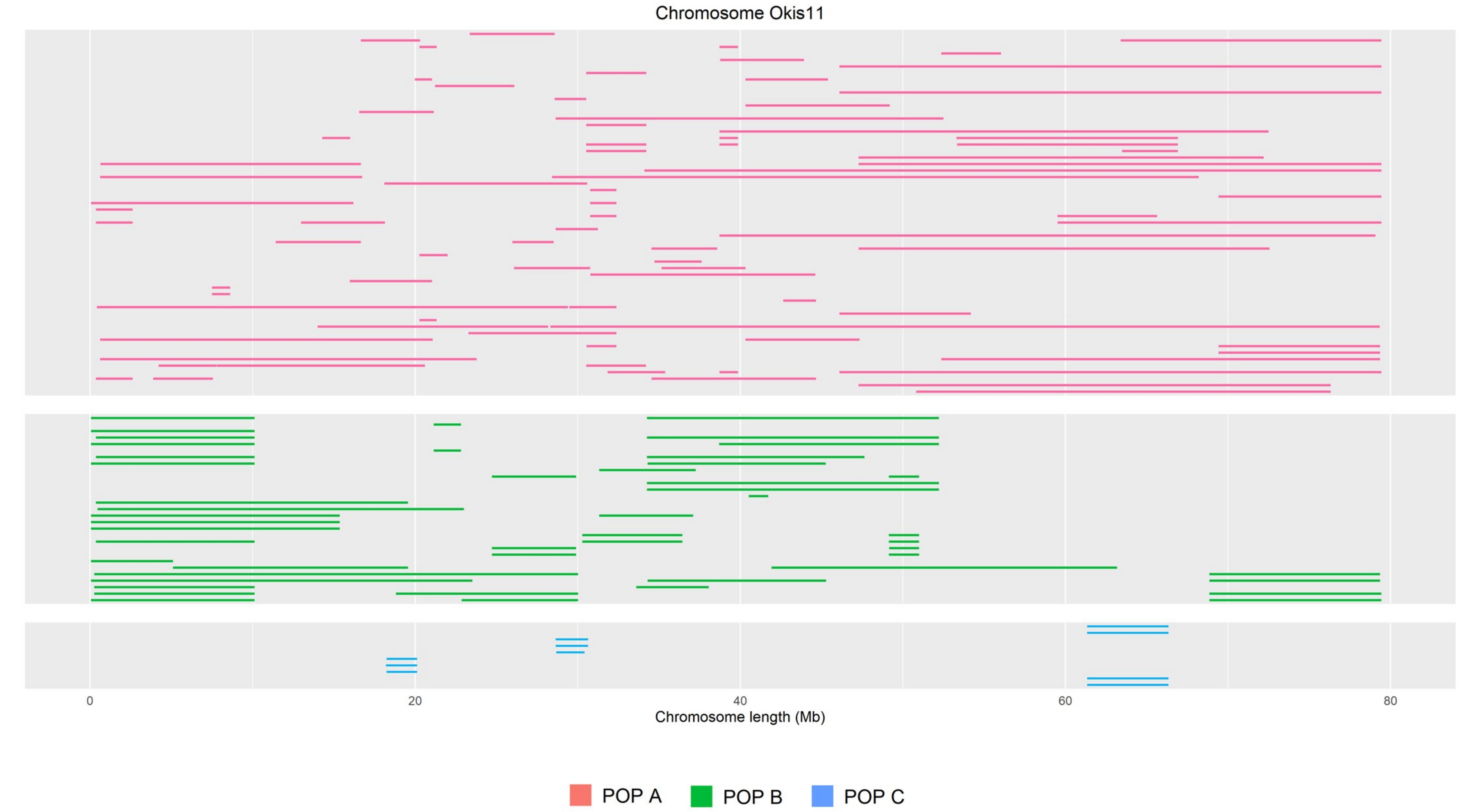
Chromosome Okis11
0
20
40
60
80
Chromosome length (Mb)
POP A
POP B
POP C

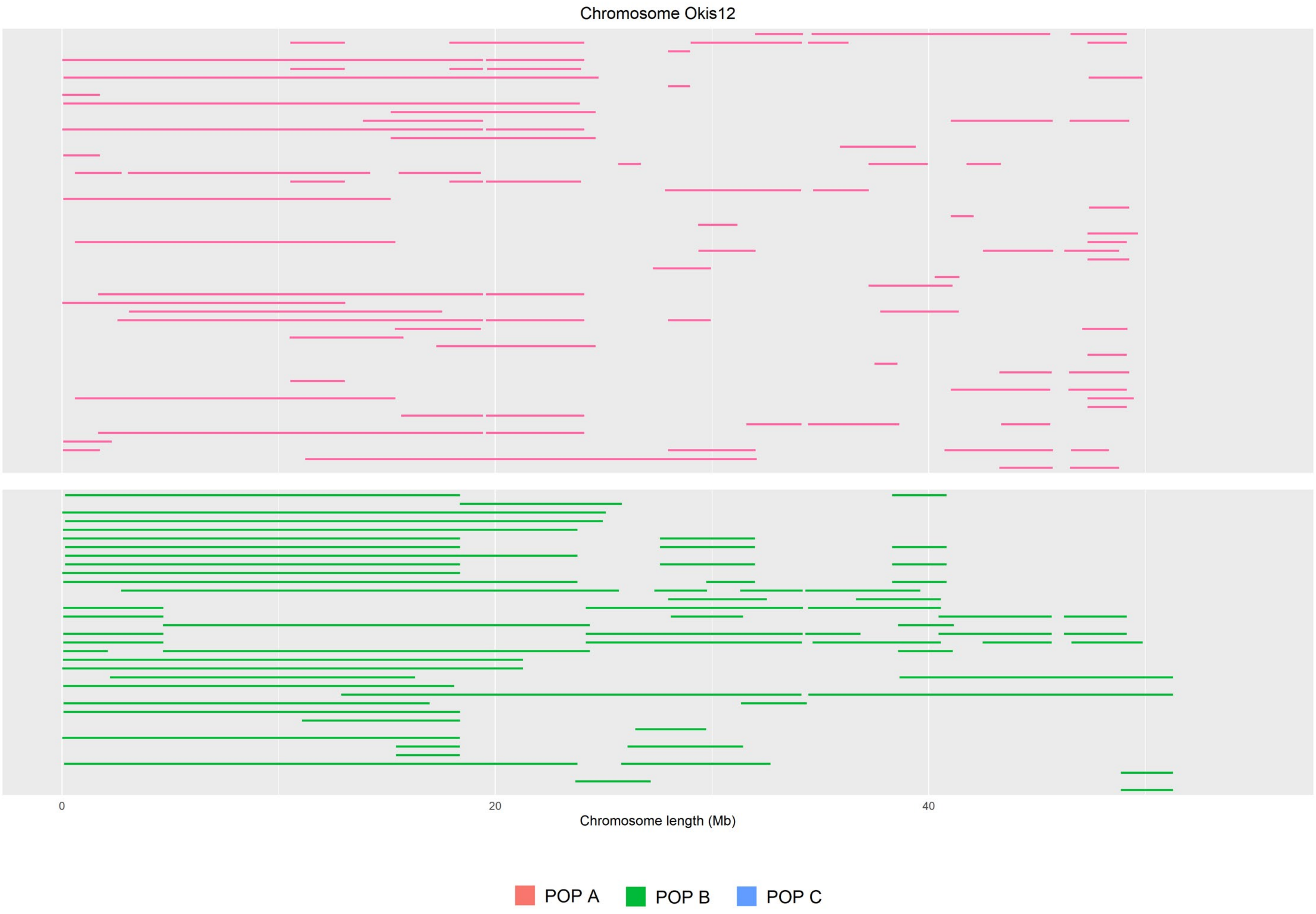

Chromosome Okis12
0
20
40
Chromosome length (Mb)
POP A
POP B
POP C

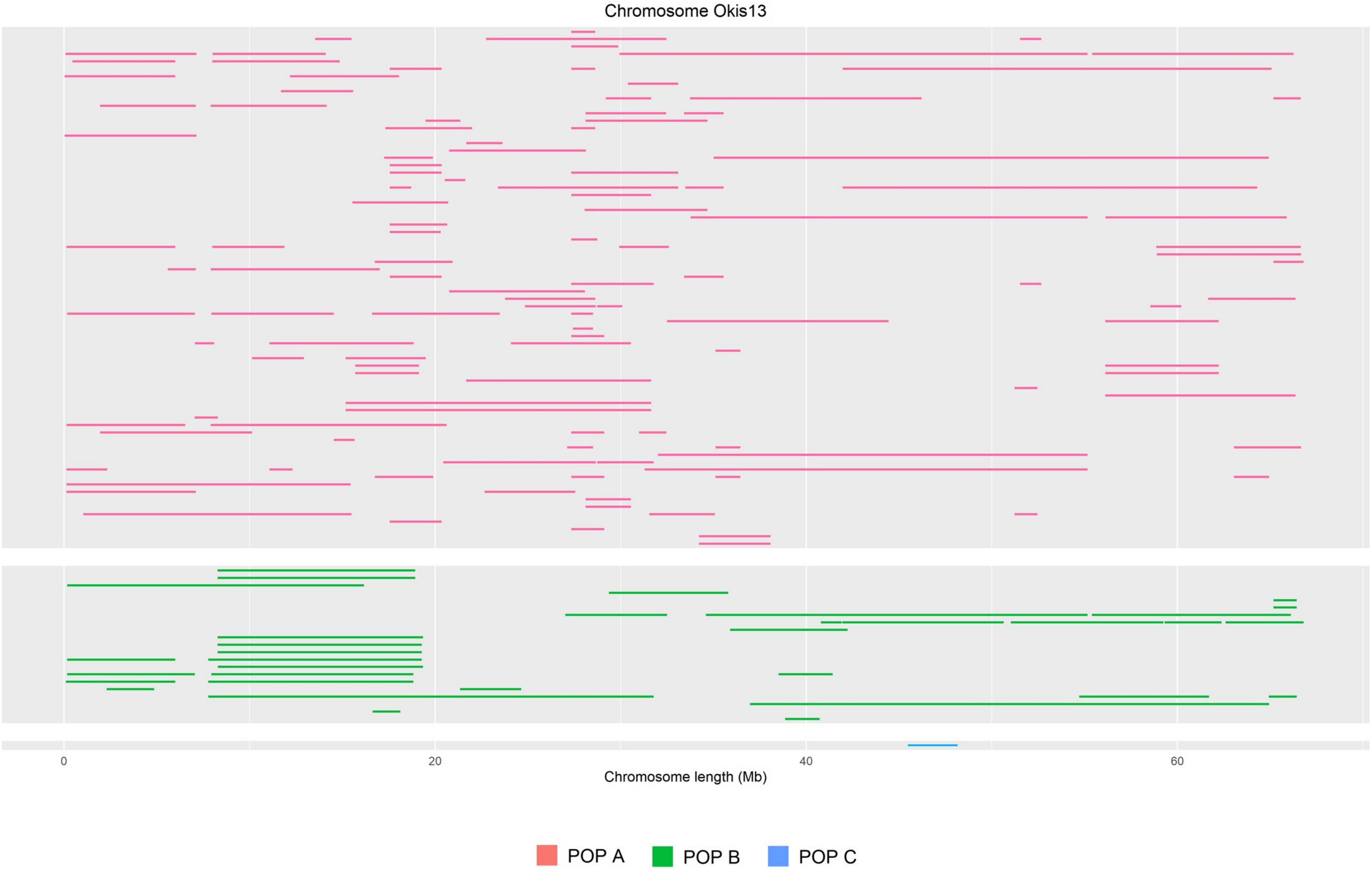

Chromosome Okis13
0
20
40
60
Chromosome length (Mb)
POP A
POP B
POP C

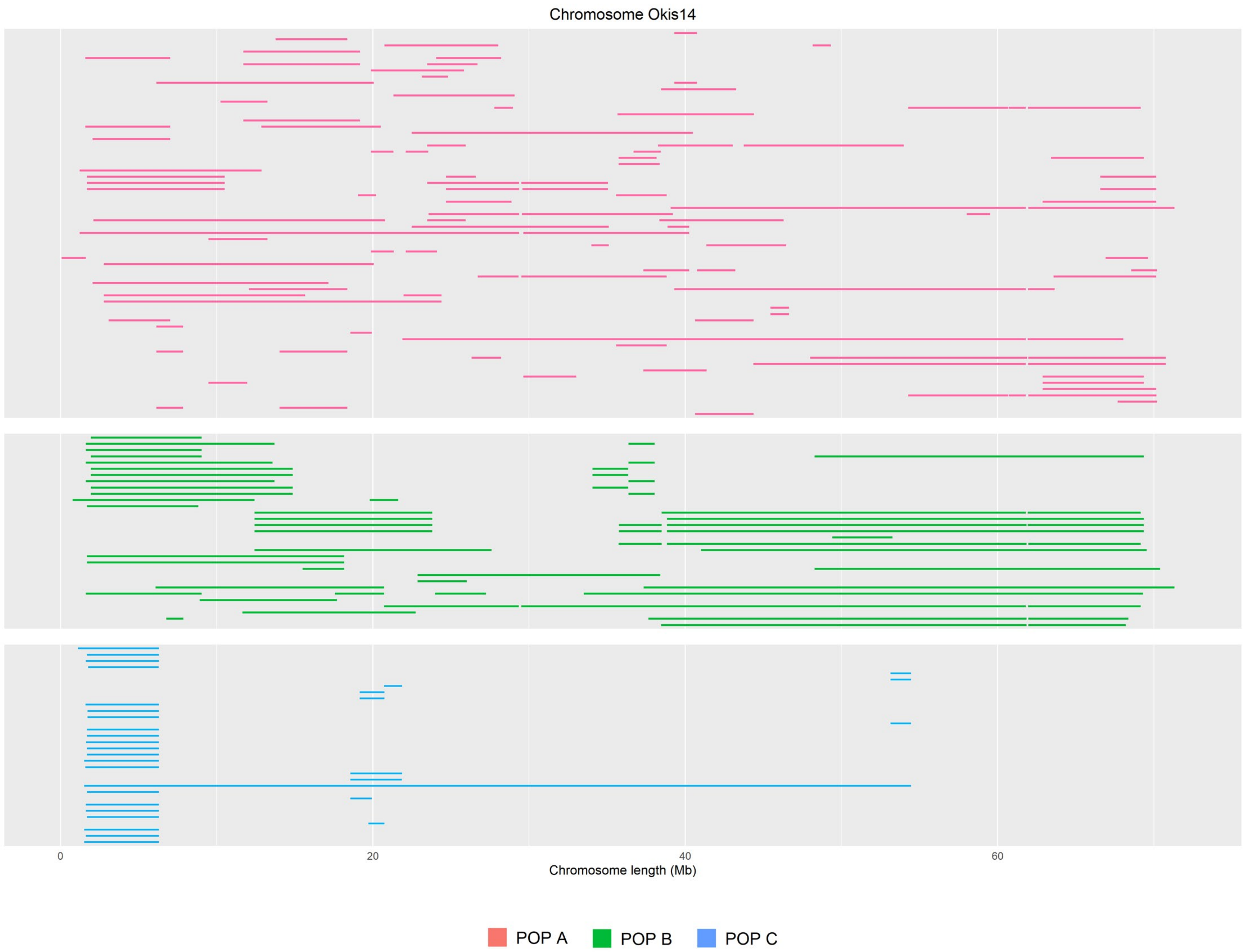

Chromosome Okis14
0
20
40
60
Chromosome length (Mb)
POP A
POP B
POP C

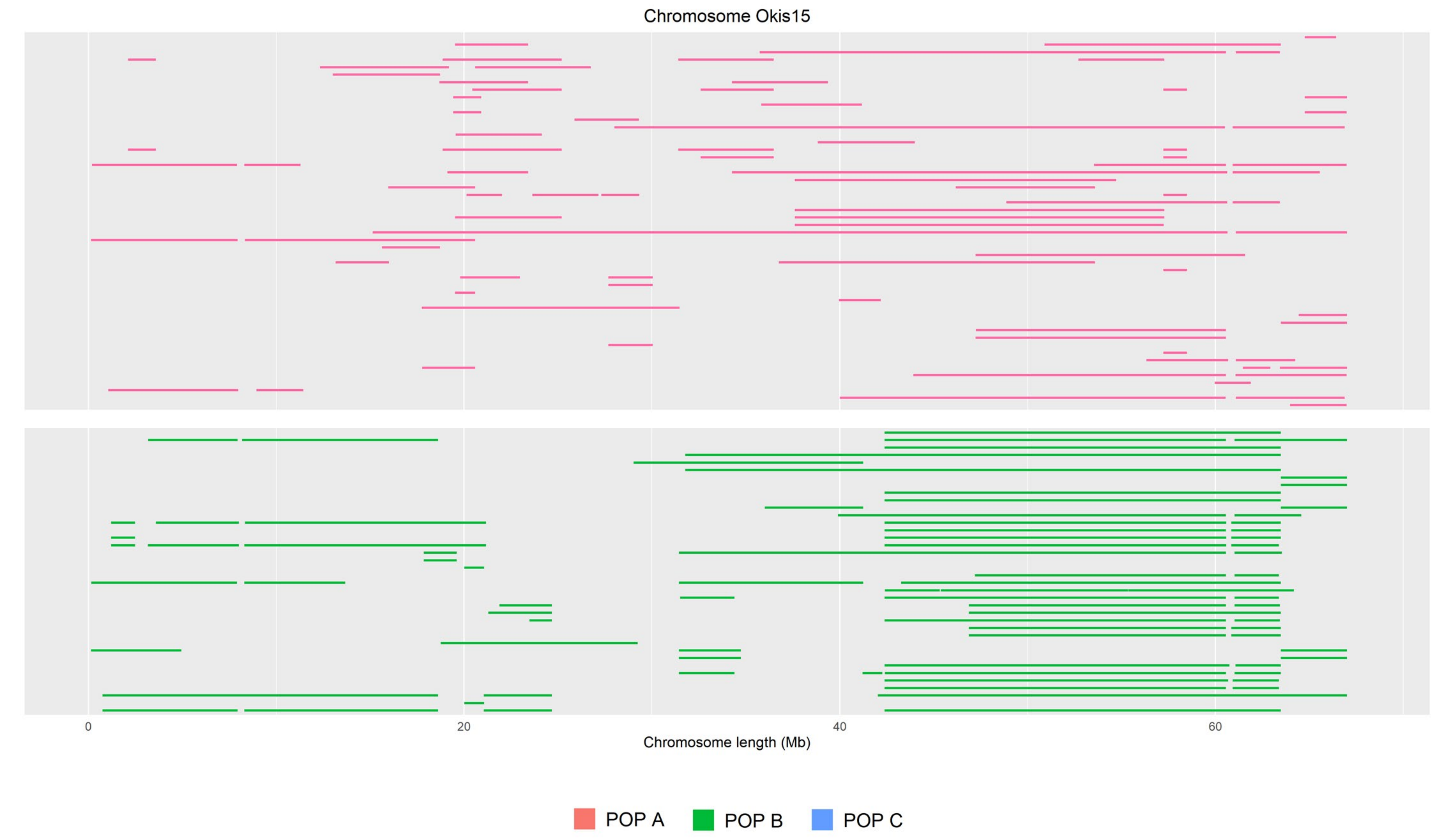

Chromosome Okis15
0
20
40
60
Chromosome length (Mb)
POP A
POP B
POP C

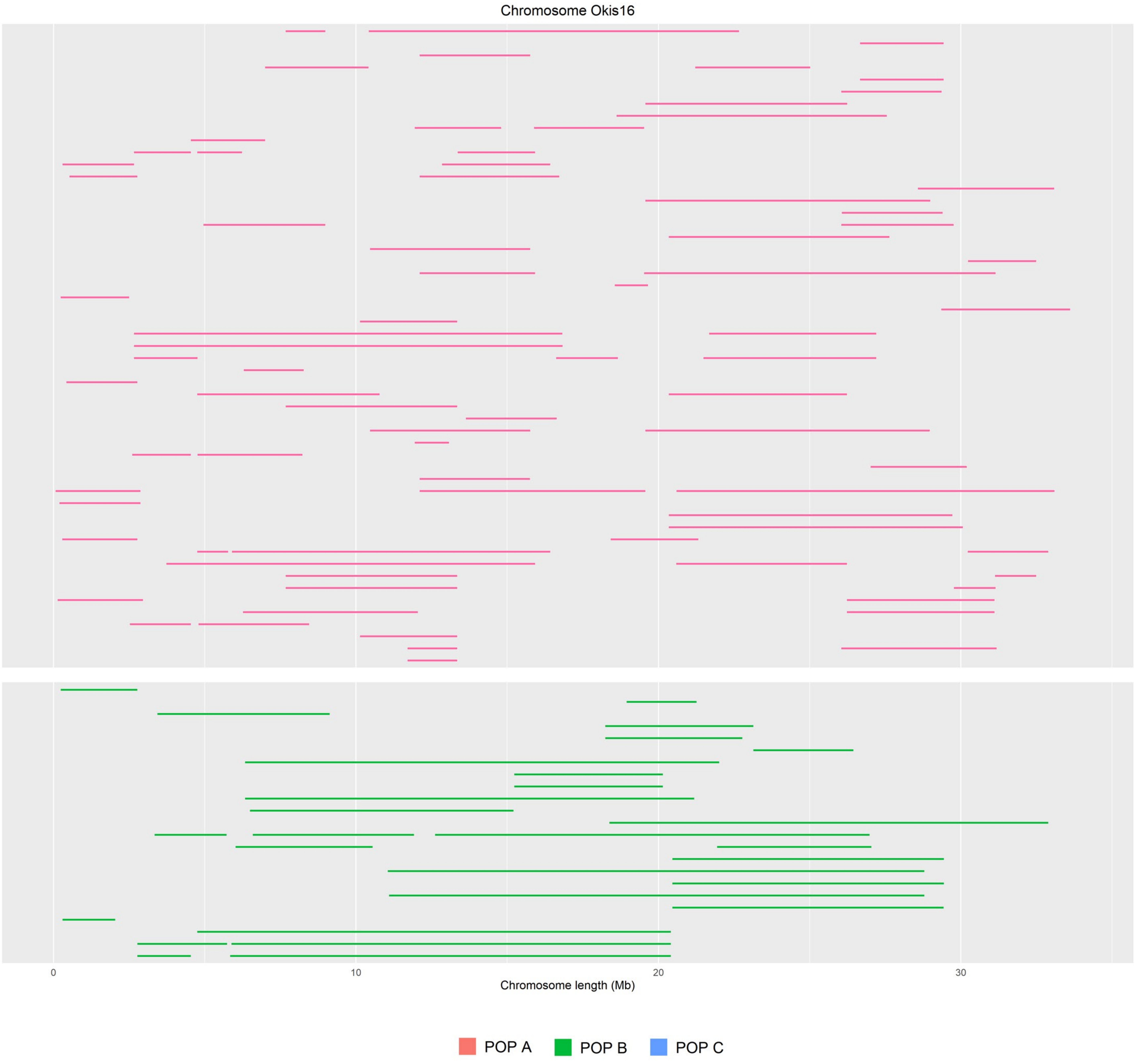
Chromosome Okis16
0
10
20
30
Chromosome length (Mb)
POP A
POP B
POP C

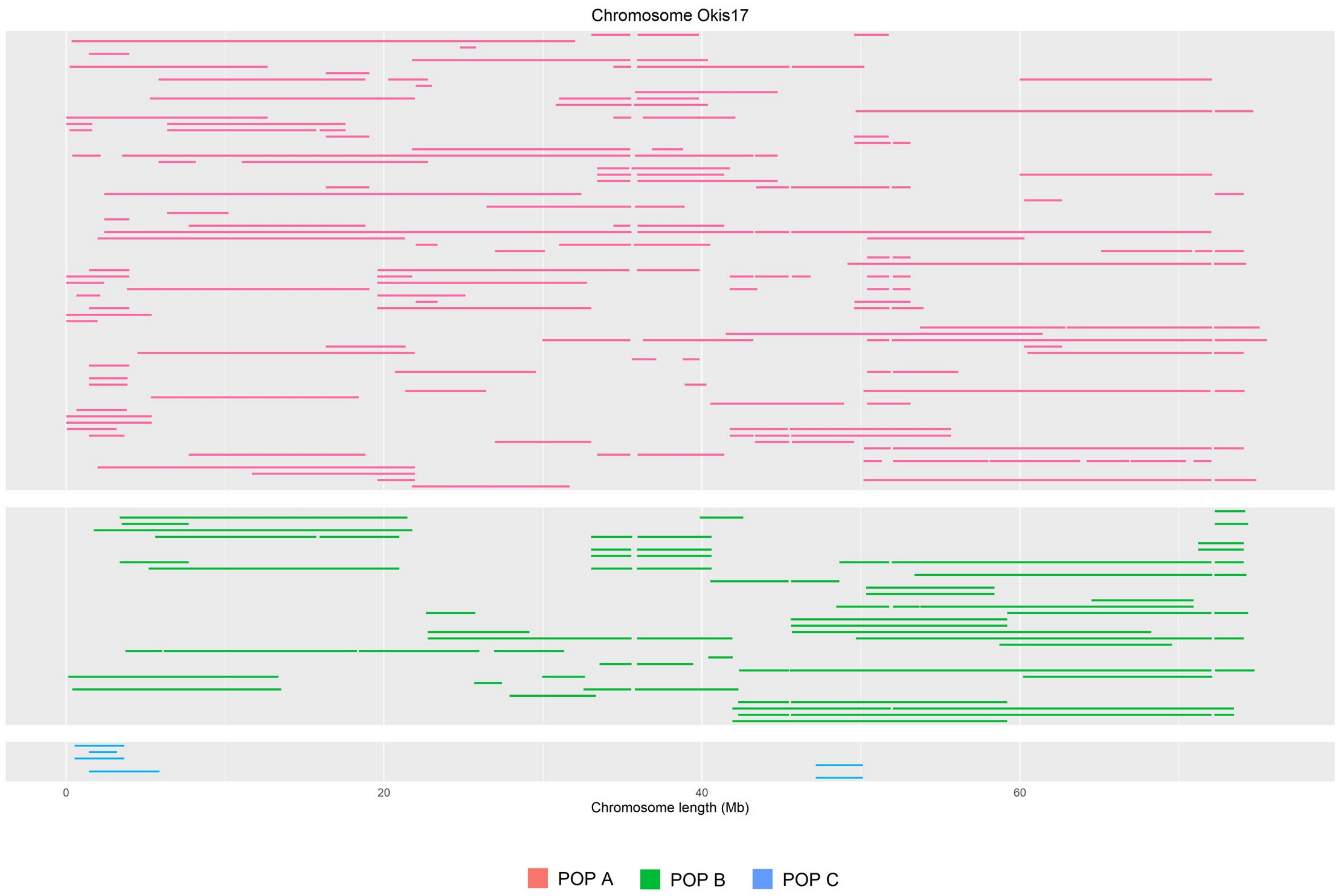

Chromosome Okis17
0
20
40
60
Chromosome length (Mb)
POP A
POP B
POP C

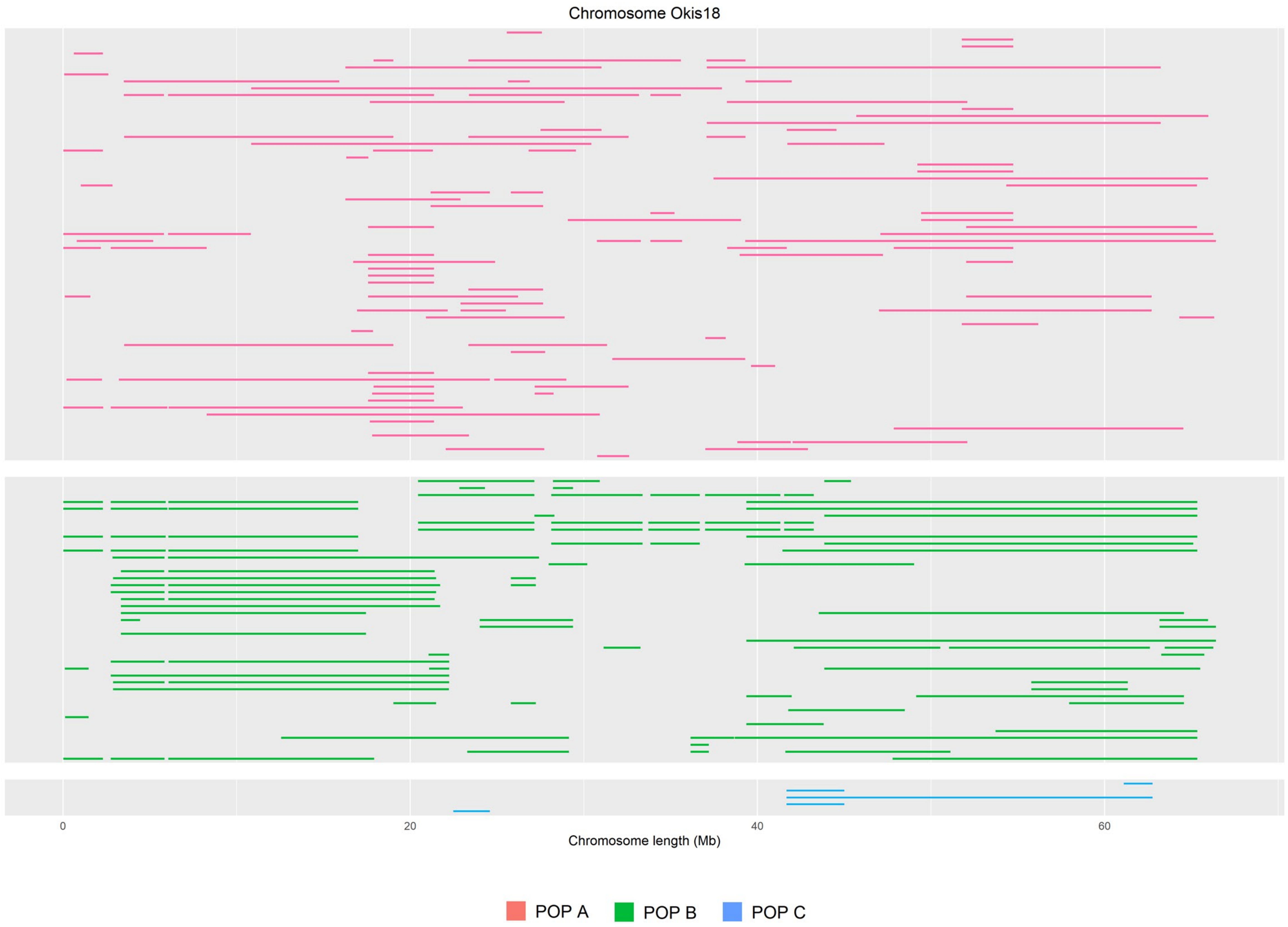
Chromosome Okis18
0
20
40
60
Chromosome length (Mb)
POP A
POP B
POP C

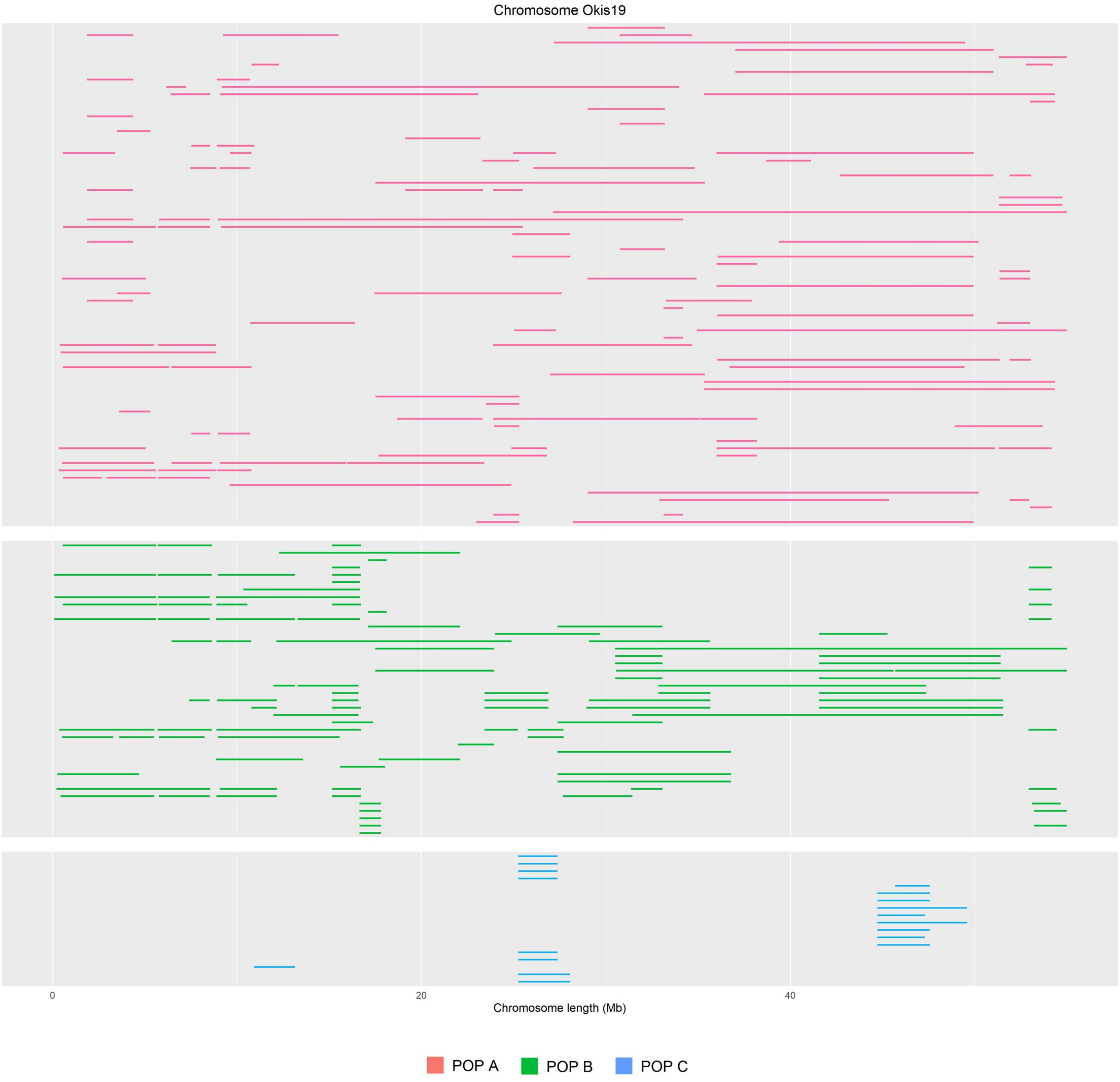

Chromosome Okis19
0
20
40
Chromosome length (Mb)
POP A
POP B
POP C

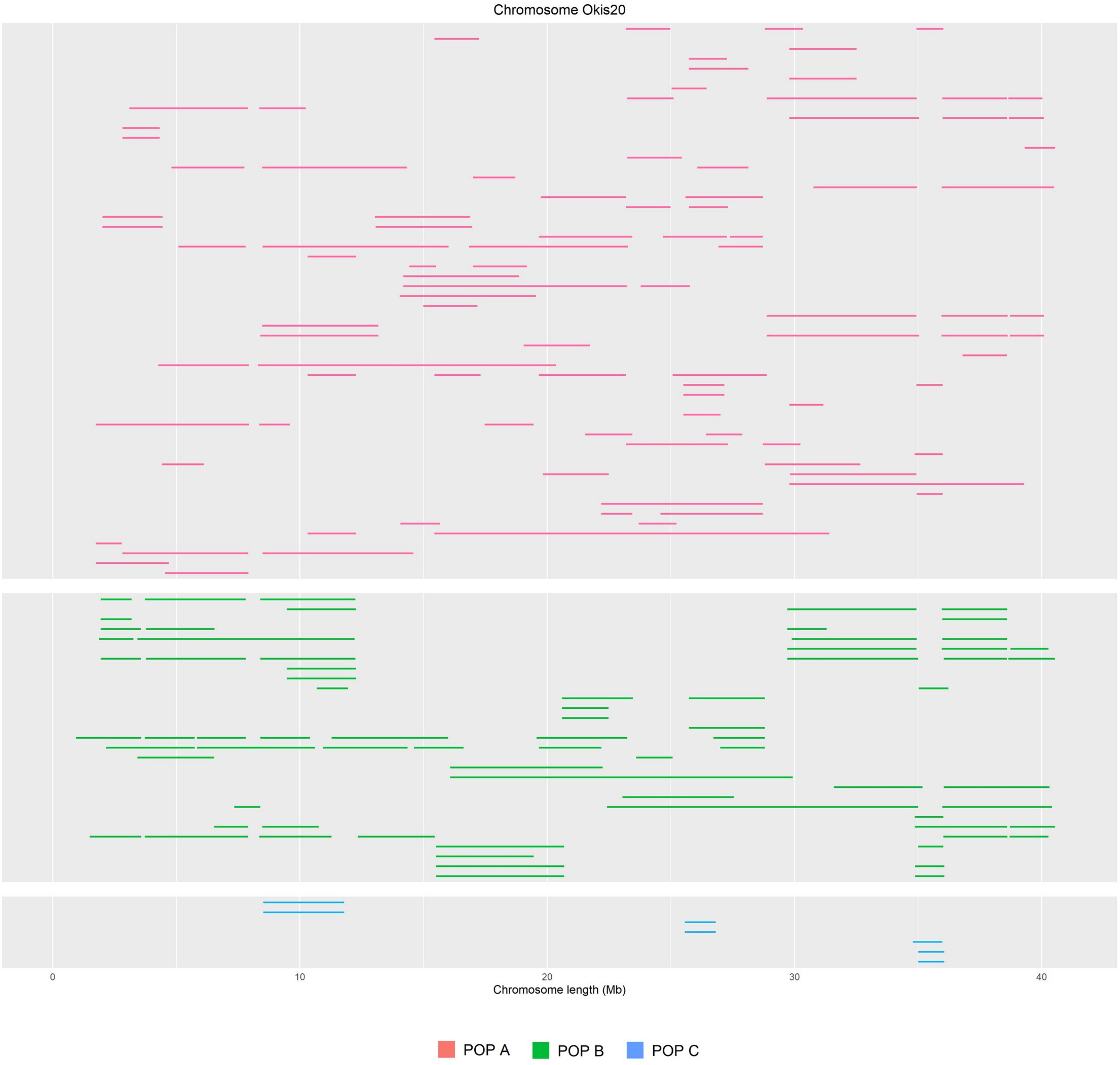

Chromosome Okis20
0
10
20
30
40
Chromosome length (Mb)
POP A
POP B
POP C

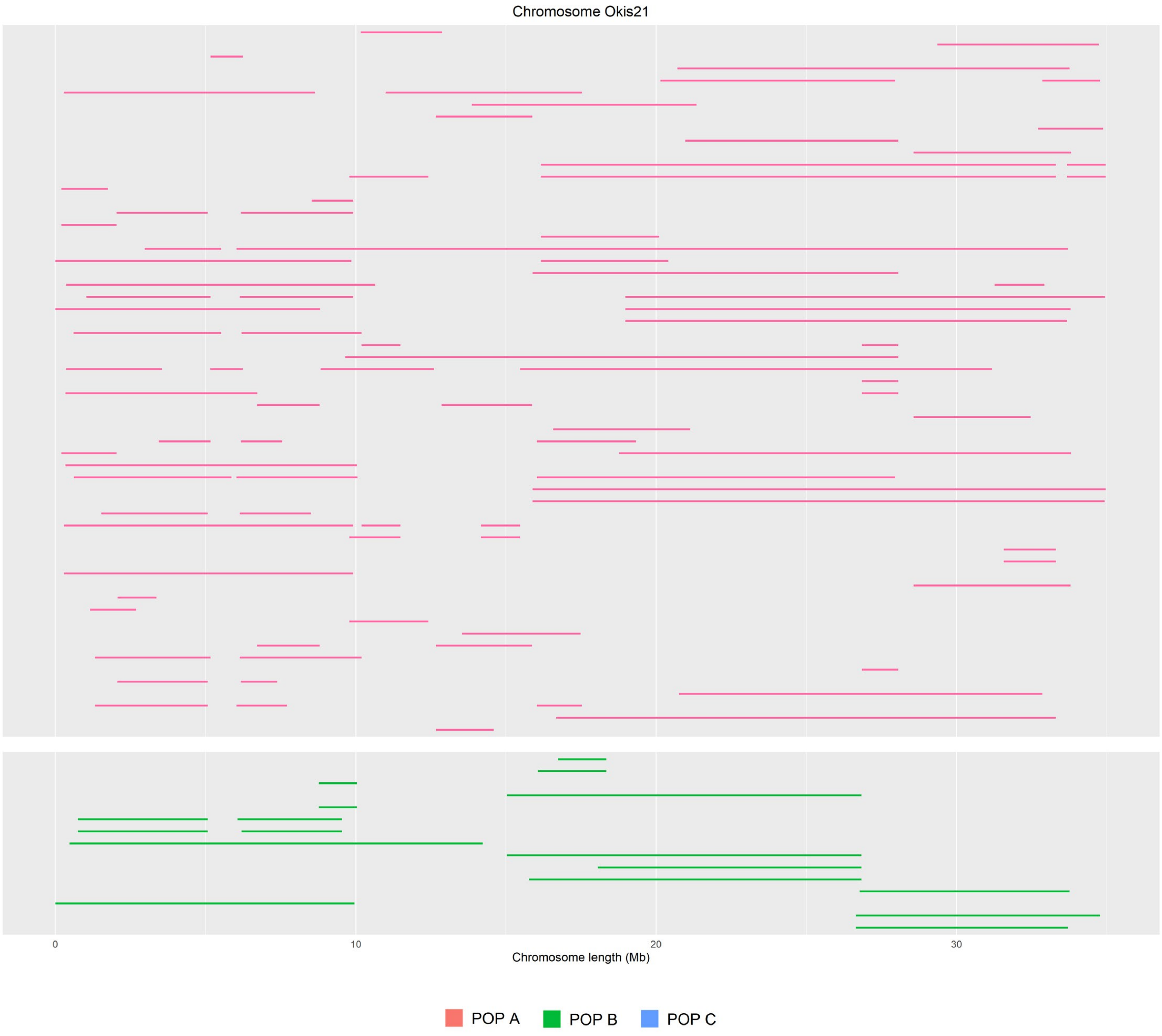

Chromosome Okis21
0
10
20
30
Chromosome length (Mb)
POP A
POP B
POP C

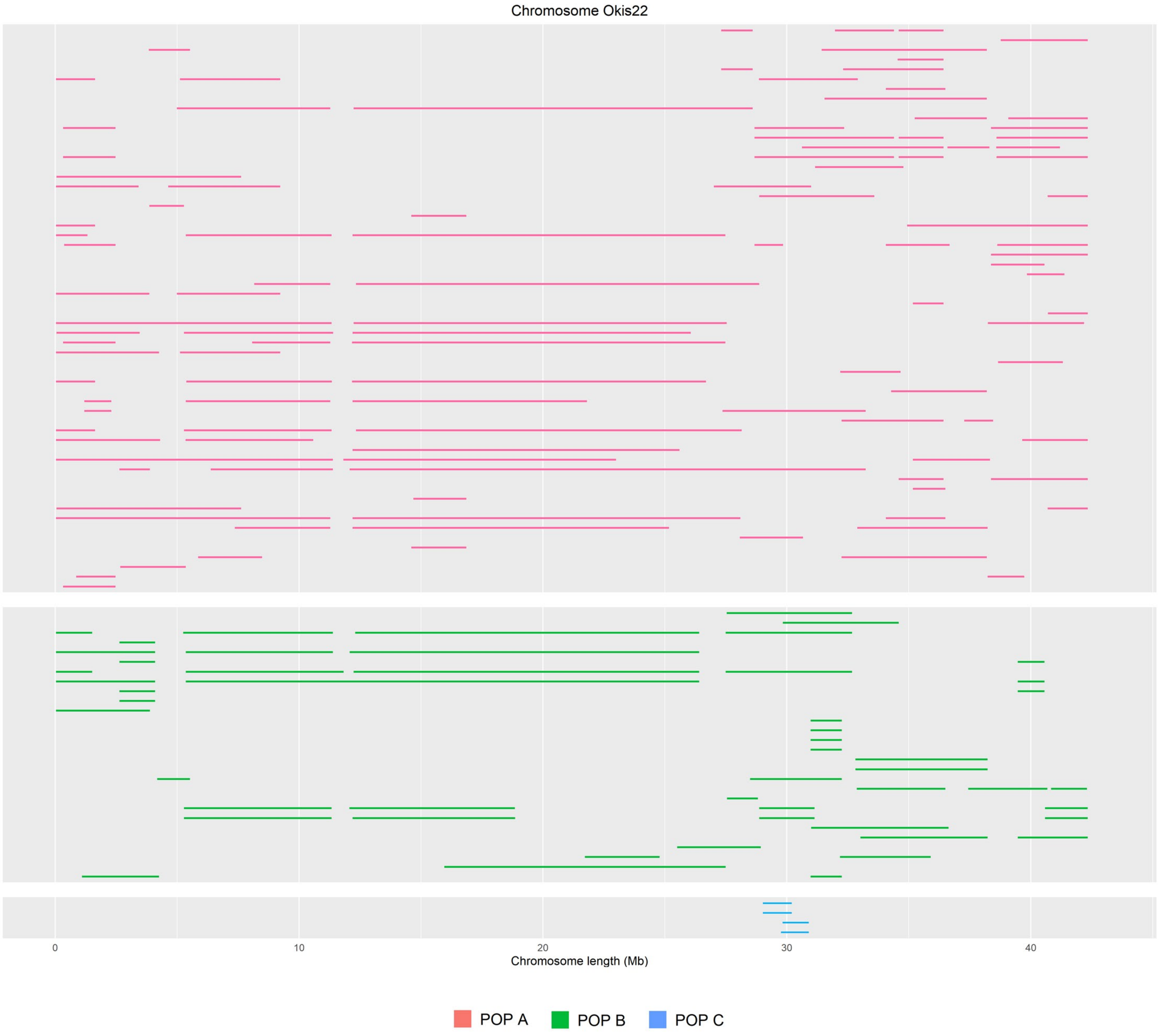

Chromosome Okis22
0
10
20
30
40
Chromosome length (Mb)
POP A
POP B
POP C

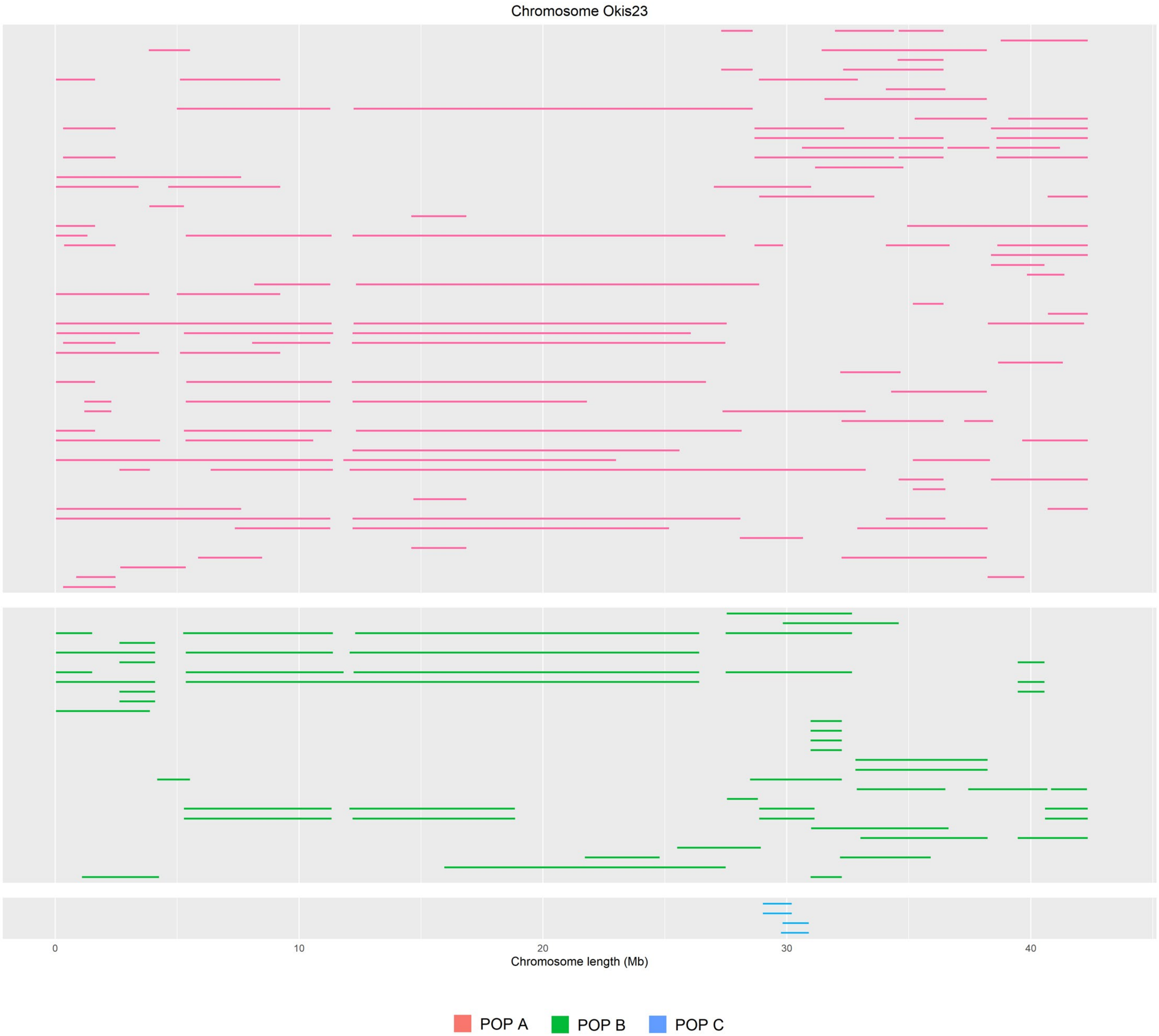
Chromosome Okis23
0
10
20
30
40
Chromosome length (Mb)
POP A
POP B
POP C

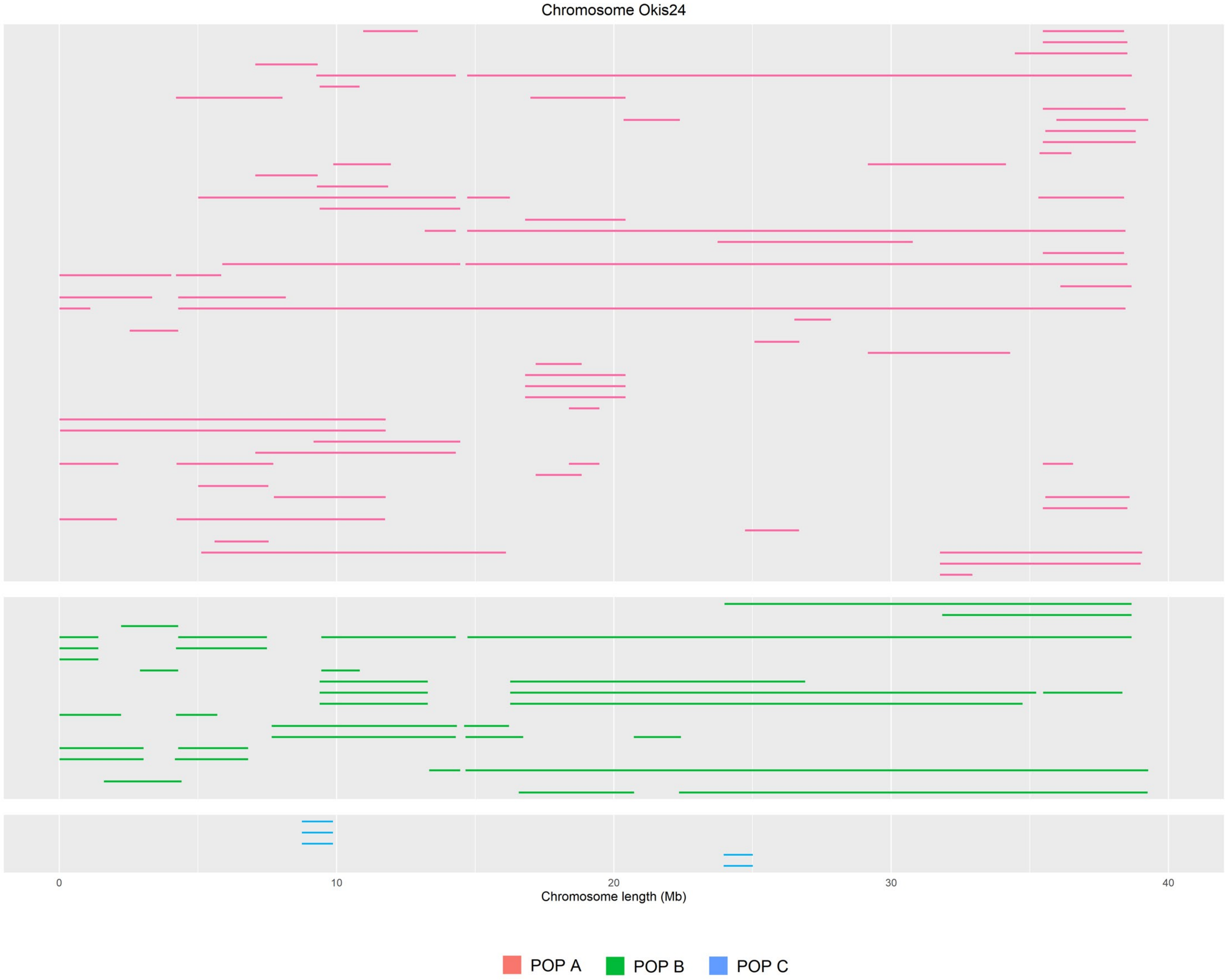

Chromosome Okis24
0
10
20
30
40
Chromosome length (Mb)
POP A
POP B
POP C

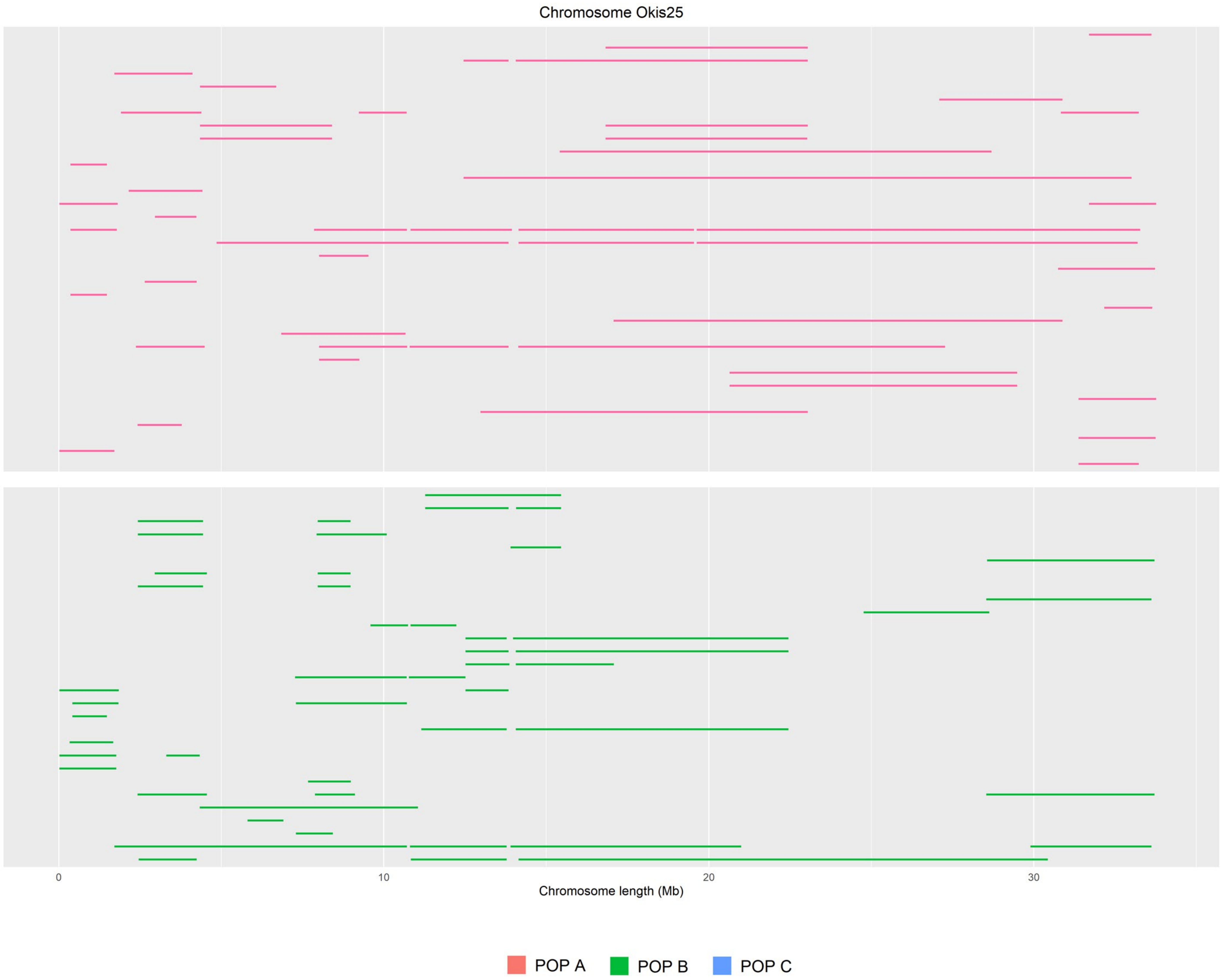

Chromosome Okis25
0
10
20
30
Chromosome length (Mb)
POP A
POP B
POP C

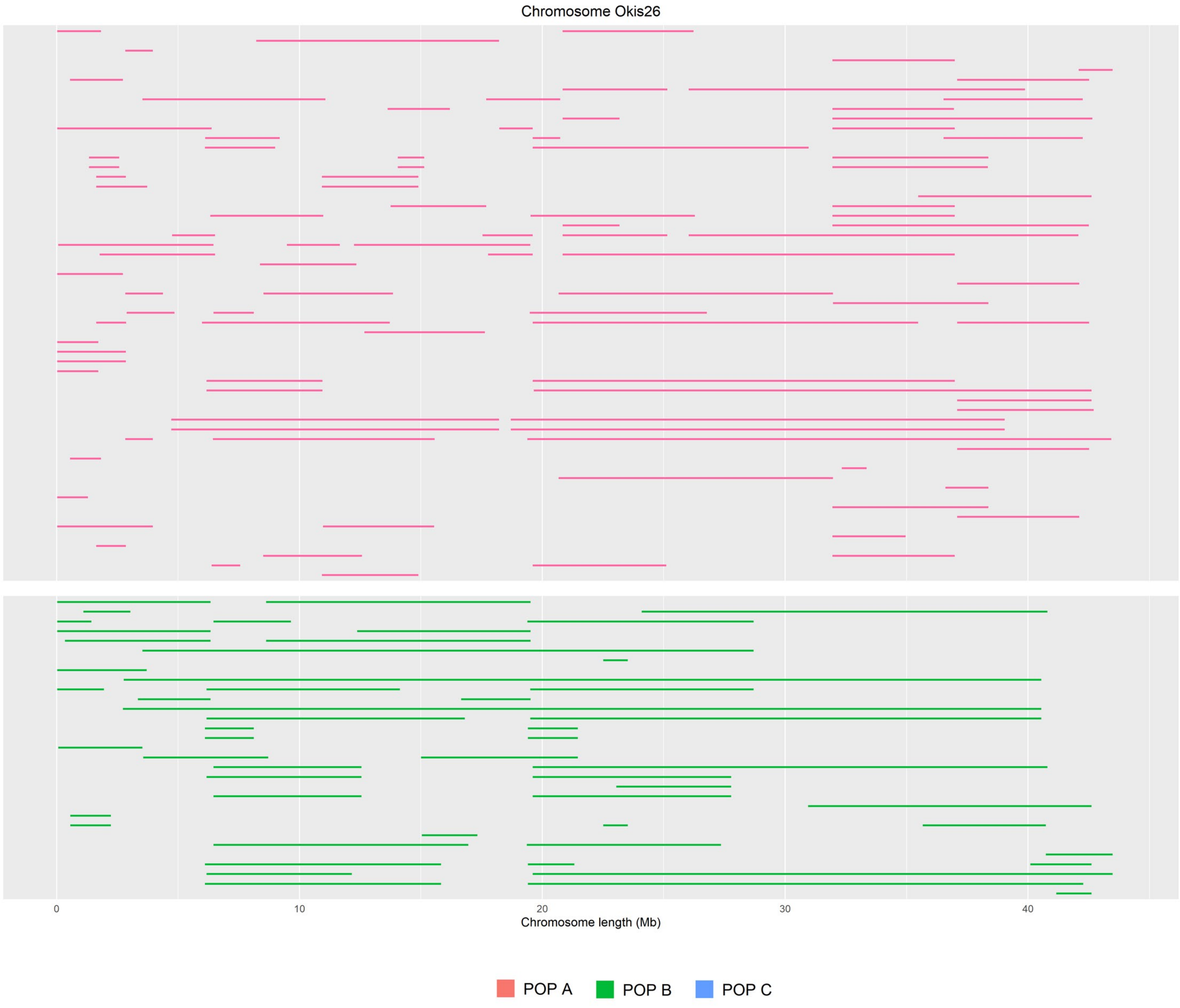
Chromosome Okis26
0
10
20
30
40
Chromosome length (Mb)
POP A
POP B
POP C

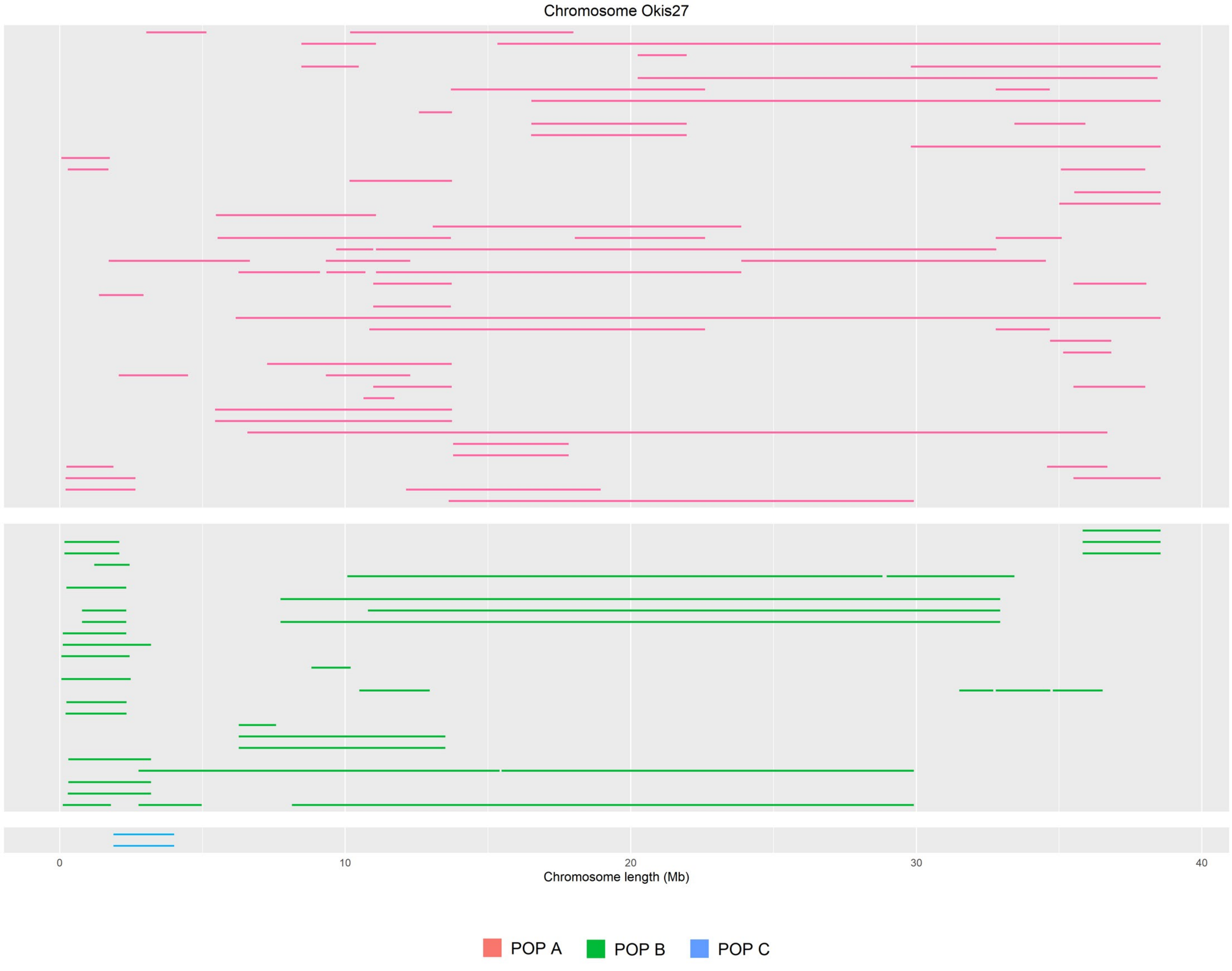

Chromosome Okis27
0
10
20
30
40
Chromosome length (Mb)
POP A
POP B
POP C

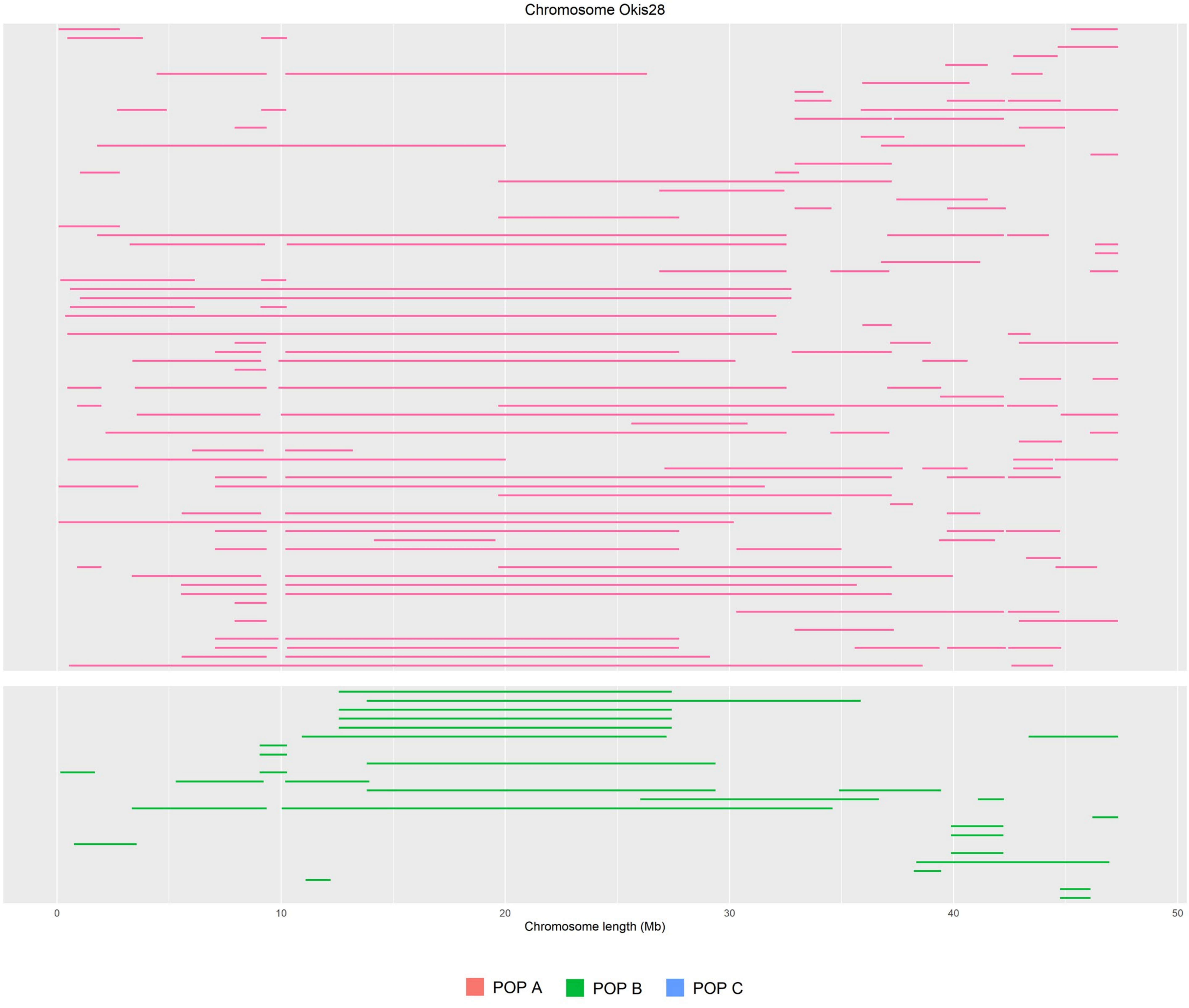

Chromosome Okis28
0
10
20
30
40
50
Chromosome length (Mb)
POP A
POP B
POP C

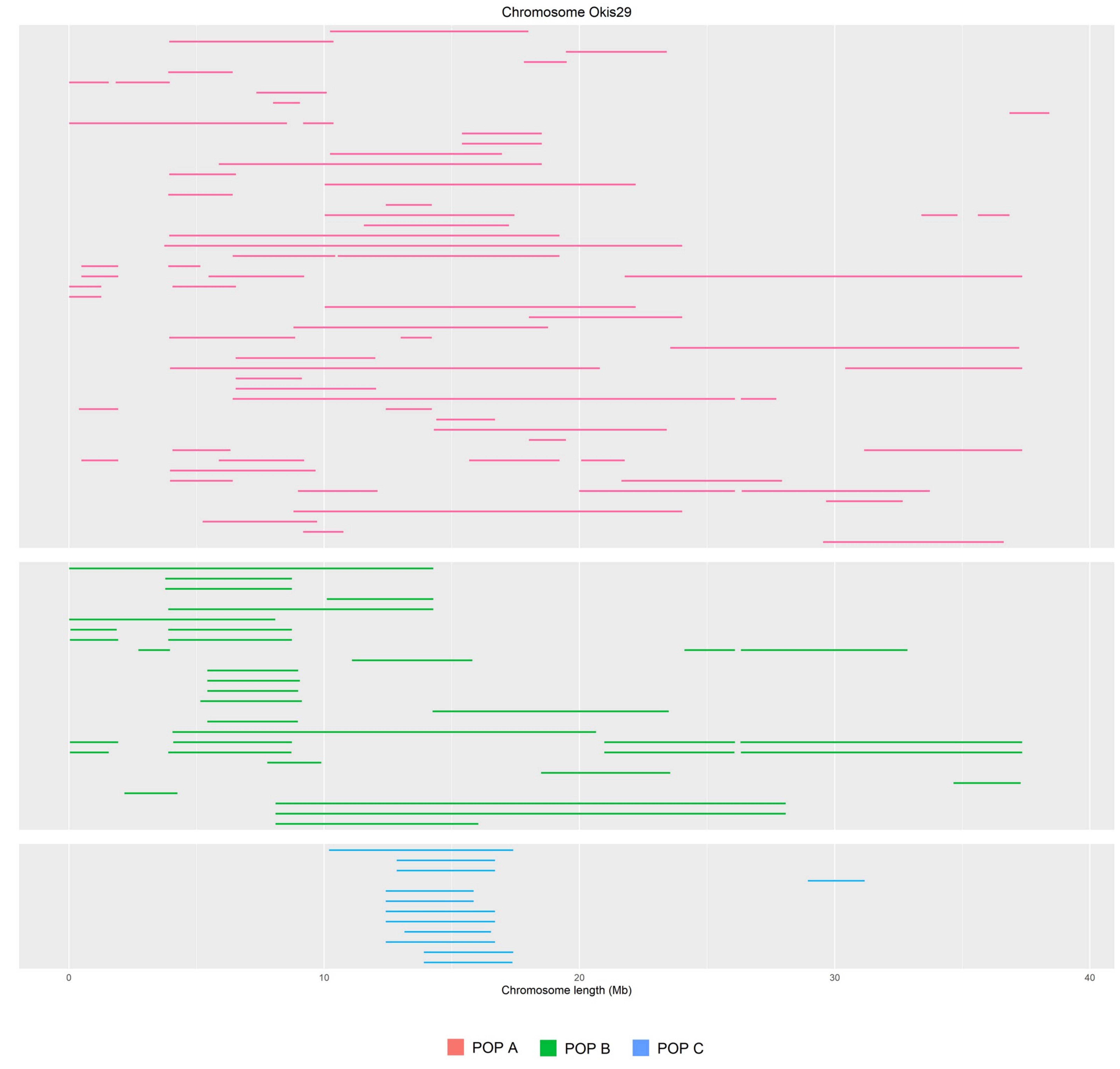

Chromosome Okis29
0
10
20
30
40
Chromosome length (Mb)
POP A
POP B
POP C

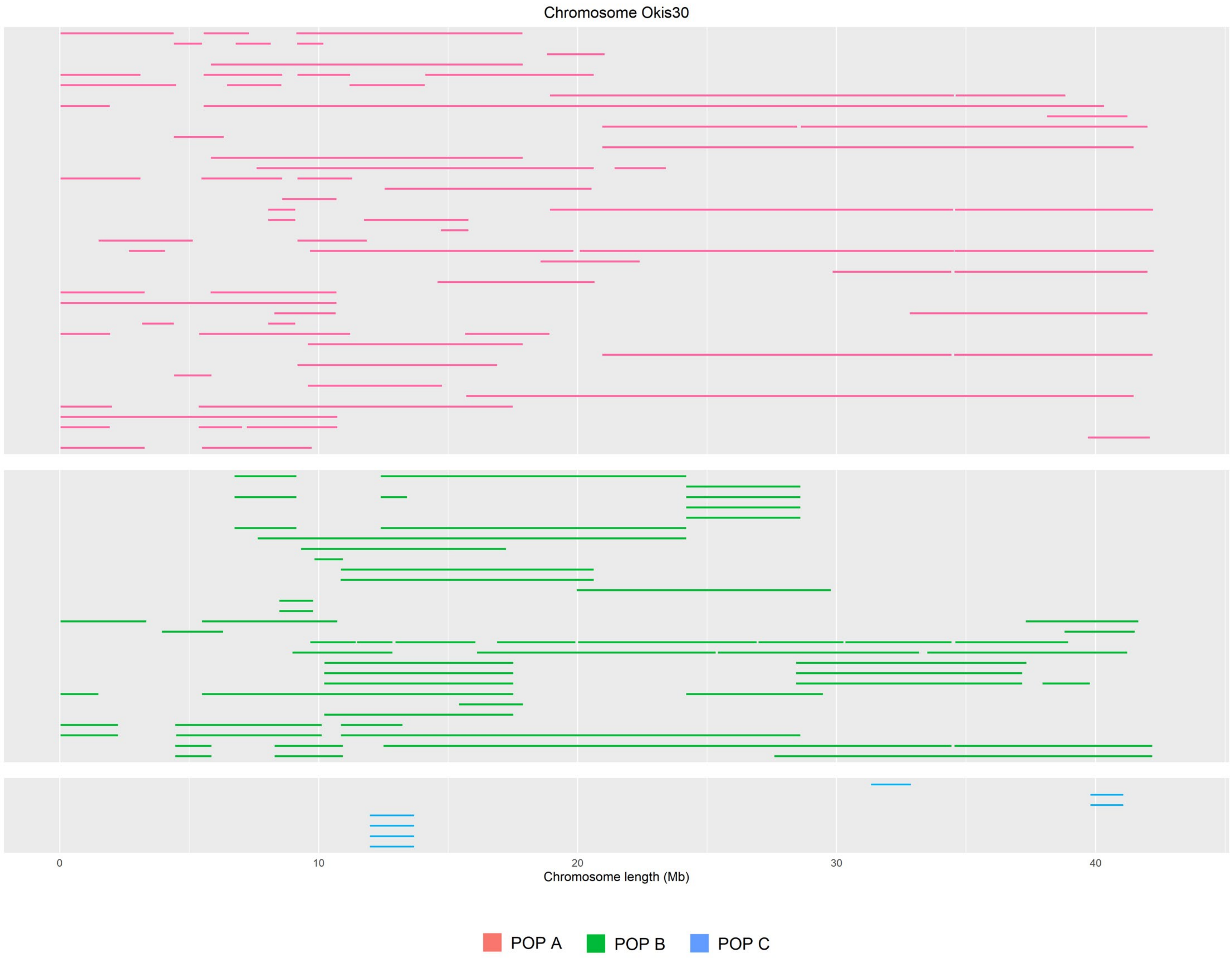

Chromosome Okis30
0
10
20
30
40
Chromosome length (Mb)
POP A
POP B
POP C
